# Chronic stress exposure alters the gut barrier: sex-specific effects on microbiota and jejunum tight junctions

**DOI:** 10.1101/2022.09.13.507821

**Authors:** Ellen Doney, Laurence Dion-Albert, Francois Coulombe-Rozon, Natasha Osborne, Renaud Bernatchez, Sam E.J. Paton, Fernanda Neutzling Kaufmann, Roseline Olory Agomma, José L. Solano, Raphael Gaumond, Katarzyna A. Dudek, Joanna Kasia Szyszkowicz, Signature Consortium, Manon Lebel, Alain Doyen, Audrey Durand, Flavie Lavoie-Cardinal, Marie-Claude Audet, Caroline Menard

## Abstract

Major depressive disorder (MDD) is the leading cause of disability worldwide. However, 30-50% of patients are unresponsive to commonly prescribed antidepressants, highlighting untapped causal biological mechanisms. Dysfunction in the microbiota-gut-brain axis, the bidirectional communications between the central nervous system and gastrointestinal tract that are modulated by gut microorganisms, has been implicated in MDD pathogenesis. Exposure to chronic stress disrupts blood-brain barrier integrity, still, little is known about intestinal barrier function in these conditions particularly for the small intestine where most food and drug absorption takes place. Thus, here we investigate how chronic social or variable stress, two mouse models of depression, impact the jejunum (JEJ) intestinal barrier in males and females. Mice were subjected to stress paradigms followed by analysis of gene expression profiles of intestinal barrier-related targets, fecal microbial composition, and blood-based markers. Altered microbial populations as well as changes in gene expression of JEJ tight junctions were observed depending on the type and duration of stress, with sex-specific effects. We took advantage of machine learning to characterize in detail morphological tight junction properties identifying a cluster of ruffled junctions in stressed animals. Junctional ruffling is associated with inflammation, so we evaluated if LPS injection recapitulates stress-induced changes in the JEJ and observed profound sex differences. Finally, LPS-binding protein (LBP), a marker of gut barrier leakiness, was associated with stress vulnerability in mice and translational value was confirmed on blood samples from women with MDD. Our results provide evidence that chronic stress disrupts intestinal barrier homeostasis in conjunction with the manifestation of depressive-like behaviors in a sex-specific manner in mice and possibly, human depression.

## Introduction

Major depressive disorder (MDD) is currently the most prevalent mood disorder and the leading cause of disability worldwide^1-3^. Depressive disorders fall along a range of severities from a milder, persistent dysthymia to severe MDD. Core symptoms include low mood, irritability, anhedonia, apathy, difficulty concentrating, disrupted appetite and sleep^1-3^. Still, this disorder is highly heterogeneous and the experience, expression of symptoms and response to treatment varies highly across individuals^1,3^. Maladaptive central and peripheral inflammatory responses are associated with mood disorders and have received increasing attention in psychiatry^4-6^. Indeed, administration of pro-inflammatory cytokines or endotoxins is sufficient to induce behavioral symptoms associated with depression and treatment-resistant individuals with MDD are characterized by elevated levels of circulating cytokines^3-6^. Aligning with the neuroimmune hypothesis of depression, MDD has a high comorbidity with inflammatory bowel diseases such as Crohn’s disease and ulcerative colitis^7,8^, suggesting that inflammation-driven gut barrier dysfunction may affect emotion regulation and vice versa^1^. In gastrointestinal disorders, elevated pro-inflammatory cytokines promote increased permeability in the intestinal tract by suppressing tight junction mediated barrier function^9^. Shared profiles of upregulated pro-inflammatory cytokines in the blood, such as interleukin-1 beta, tumor necrosis factor-alpha, interleukin-6 and interferon-gamma, occur in gastrointestinal disorders and MDD, which could be related to increased intestinal permeability^1,7^. Stress has been linked to the deterioration of the intestinal barrier via alterations of gut–brain signaling^10,11^. We recently showed that it can alter blood-brain barrier (BBB) integrity in a sex-specific manner through loss of the tight junction protein Claudin-5 (Cldn5), leading to the development of anxiety- and depression-like behaviors^12-15^. To our knowledge, the impact of stress exposure on gut barrier integrity and assessment of sex differences has yet to be determined.

Women are twice more likely than men to be diagnosed with MDD^16^. Symptomatology and treatment responses also differ between sexes^17-19^. As chronic stress is the main environmental risk factor for MDD^20^, it is commonly used in male and female rodents to induce depression-like behaviors and investigate underlying biology. Chronic social defeat stress (CSDS) is a mouse model of depression based on social dominance, which produces two distinct phenotypes of stress response: stress-susceptible (SS) and resilient (RES) mice^21-23^. The SS subgroup display distinct behavioral changes reminiscent of depressive symptoms in humans with increased social avoidance, anxiety, anhedonia, despair, body weight changes, metabolic disturbances, and corticosterone reactivity^21-23^. Furthermore, loss of BBB integrity occurs only in the brain of SS, but not RES, mice^12,14^. Another leading stress paradigm is the chronic variable stress (CVS) model, during which mice are exposed to a repetitive sequence of three stressors, namely tube restraint, tail suspension, and foot shocks. Each stressor endures about 1 hour daily and lasts from 6 days to several weeks^24,25^. CVS also induces a pro-inflammatory immune profile like that produced by CSDS^26^. In this paradigm, females and males develop depression-like behaviors at different time points making it a strong model for investigating sex differences^4,25^.

The gut barrier is formed by the mucus layer, the epithelia, and a connective tissue layer, known as the lamina propria^27^. The epithelial cell monolayer faces the luminal side, interacting with the environment and regulating absorption and secretion. In the small intestine, macro-structures of the epithelial surface consist of elongated villi that protrude into the lumen and crypts of proliferating and regenerating cells at the base between them^27^. The epithelium provides a dynamic and semi-permeable barrier with tight junction complexes, linking adjacent cells, mediating the extent of the various functions. Combined, junction complexes and the overlying mucus layer, maintain a healthy functional barrier which allows the passage of nutrients, water, and ions, but limits entry of pathogens and bacterial toxins from the lumen^27,28^. The intestinal barrier is at the forefront of immune-environment interactions where the specialized cells play a critical role in maintaining health through diverse mechanisms^1,29^. Permeability of the epithelial layer may increase interaction of antigens with immune cells, propagating a pro-inflammatory response^1^. Microbial translocation from the intestinal lumen into the systemic circulation in the absence of acute infection is proposed as a mechanism behind the chronic inflammation in MDD^30-32^. Indeed, the passage of bacterial products and immune factors as indirect measures of bacterial translocation has been reported in MDD^30-32^.

To assess the impact of stress exposure on gut barrier integrity and particularly the small intestine which remains understudied, we used complementary mouse models of MDD and combined behavioral, molecular, morphological, and pharmacological experiments with blood-based assays. Our results provide characterization, in a sex-specific manner, of stress-induced changes in the jejunum (JEJ) following exposure to social or variable stressors. We developed tools and algorithms to analyze in detail tight junction morphological changes and identified circulating LPS-binding protein (LBP) as a gut leakiness potential biomarker that could help better diagnose and inform treatment strategies for mood disorders.

## Results

### Chronic social stress alters intestinal tight junction expression with sex-specific effects

10-day CSDS exposure induced expression of SS and RES phenotypes based on the social interaction (SI) test (**Fig.1A**). Among the 29 male mice subjected to social defeat, 16 had SI ratio <1 and were classified as SS (55.2%) and 13 had SI ratios of ≥1 and were classified RES (44.8%) (**Fig.1B**, left panel, SI ratio: *p<0.0001*). Time spent in the corners when the aggressor (AGG) was present is increased for SS mice (**Fig.1B**, middle panel, *p<0.0004*) while total distance traveled was similar between groups (**Fig.1B**, right panel and **Supp. Fig.1A-B**). 24h after the SI test, tissue was collected and mRNA transcriptional profiling was performed for genes related to intestinal tight junctions (*Cldn3, Cldn7, Cldn12*), tight junction-associated proteins (*Tjp1, Tjp2, Tjp3, Ocln*, MARVEL domain-containing protein 2 *[Marveld2]*), as well as proteins involved in serotonin metabolism (*Ido1* and *Ahr*) and mucus layer formation, Mucin-2 (*Muc2*) on the jejunum (JEJ) of unstressed controls (CTRL), SS and RES mice. Indeed, serotonin metabolism is altered in inflammatory bowel diseases, which are highly comorbid with MDD^7^, and the mucus layer is essential to maintain gut barrier integrity. Fold changes were positively correlated with SI ratios for *Cldn12* (*p=0.006*, r=0.44), *Tjp1* (*p=0.03, r=0.36*), *Tjp2* (*p=0.004, r=0.46*) and *Ocln* (*p<0.001, r=0.55*) (**Fig.1C**), suggesting that loss of intestinal barrier integrity may be linked to social avoidance in SS male mice. Intriguingly, *Cldn3*, an important intestinal tight junction, was upregulated after CSDS exposure (**Fig.1D**, *p=0.0002*).

**Figure 1.**
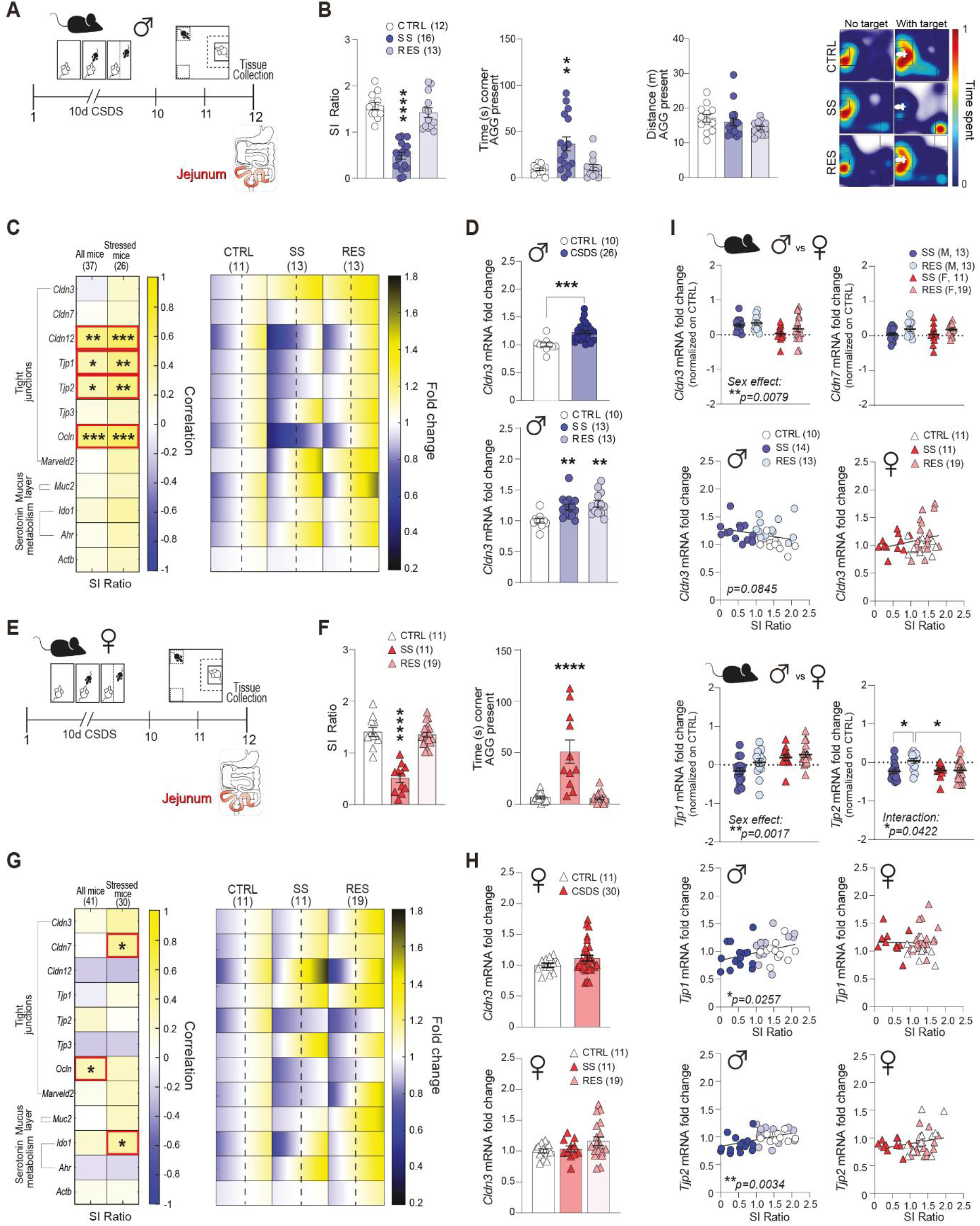
Chronic social stress alters jejunum (JEJ) tight junction expression with sex-specific effects. **A)** Timeline of the male chronic social defeat (CSDS) paradigm. **B)** Ratio of time spent interacting with novel social target decreased in stress-susceptible (SS) vs unstressed control (CTRL) and resilient (RES) male mice. Cumulative time (in seconds [s]) spent in corners with social target present was increased in SS males. Cumulative distance traveled (meters [m]) in arena with social target present was unchanged between groups. Representative heatmaps of normalized time spent in the arena during social interaction test in males. **C**) Effects of social stress on the mRNA expression of tight junction proteins in the jejunum of male mice as a function of group condition (left). Red boxes highlight genes with significant correlation with social avoidance. Quantitative PCR revealed significant changes in jejunum of SS and RES mice compared to controls for gene expression of targets related to tight junctions (right). The range of color indicates individual differences within a group; S.E.M. from the average represented by the dashed line. **D)** Significant increase in *Cldn3* for male mice was independent of phenotype group. **E)** Timeline of the female CSDS paradigm. **F)** Ratio of time spend interacting with novel social target is decreased in SS female mice. Cumulative time (in seconds [s]) spent in corners with social target present was increased in SS females. **G)** *Ocln, Cldn7* and *Ido1* expression correlated with social avoidance behaviours (left). Red boxes highlight genes with significant correlation with social avoidance. Tight junction changes as a function of phenotype in female mice (right). **H)** *Cldn3* expression is unchanged in female mice following social stress. **I**) There is an effect of Sex as a factor on *Cldn3* and *Tjp1* gene expression, and an interaction occurs between the factor Sex and behavioural phenotype on *Tjp2* expression. Data are assessed by T-tests and one-way ANOVA followed by Bonferroni’s multiple comparison test for changes between groups; two-way ANOVA followed by Bonferroni’s multiple comparison test for comparison between sexes; correlations were evaluated with Pearson’s correlation coefficient; \**p*<0.05, \*\**p*<0.01, \*\*\**p*<0.001.

We recently reported that CSDS alters BBB integrity in a sex-specific manner^12,14^, which may contribute to the sex differences observed in MDD prevalence, symptoms, and treatment response. In female mice, exposure to CSDS (**Fig.1E**) also leads to two subpopulations of SS (36.7%) and RES (63.3%) mice with SS mice displaying social avoidance (**Fig.1F**, SI ratio: *p<0.0001*; time corners: *p<0.0001* and **Supp. Fig.1C-D**) however, a correlation was noted only for *Ocln* jejunum expression with SI ratio *(p=0.04*, r=0.31) (**Fig.1G**). As for stressed mice, a significant correlation was observed for *Cldn7* (*p=0.02*, r=0.41) and *Ido1* (*p=0.04*, r=0.24) (**Fig.1G**), highlighting stress-induced sex-specific patterns despite exposure to the same paradigm. Unlike in males, *Cldn3* was not elevated in female mice after CSDS (**Fig. 1H**) leading us to explore further sex differences. In fact, *Cldn3* expression was higher in unstressed female mice when compared to their male counterparts (*Cldn3*: *p<0.0001*, **Supp. Fig.1E**) indicating baseline sex differences for the JEJ tight junctions. There was a main effect of sex on *Cldn3* (*p=0.008*) and *Tjp1* (*p=0.002*) expression in SS and RES male and female mice (**Fig.1I**). Post hoc tests confirmed that stressed female mice had lower *Cldn3* expression than males in both SS and RES groups (*p=0.0499*). Similarly, SS males had lower *Tjp1* expression than SS females (*p=0.01*) and an interaction between sex and behavioral phenotype was present for *Tjp2* expression (*p=0.04*) (**Fig.1I**). Therefore, CSDS modified JEJ tight junction expression in both male and female mice, but these changes were specific to each sex.

### Changes in jejunum tight junction expression are dependent of stress type and duration

Males and females are often the same at baseline but when exposure to a stressor occurs, an effect can be observed in only one sex or have a greater effect in one of them. An example of that is subthreshold CVS (SCVS), female mice are more vulnerable to it with exposure to only 6 days of stressors being sufficient to induce anxiety- and depression-like behaviors in females but not males^25^. Considering the sex-specific effects of 10-day exposure to CSDS (**Fig.1**), we compared the impact of stress type and duration on *Cldn3* expression, one of the most abundant tight junction proteins of the JEJ^33^. While social stress increased *Cldn3* in males (**Fig.2A**, *p=0.0002*), it remained unchanged after 6-d SCVS (**Fig.2B**, left) in line with unaltered behaviors^24^. Conversely, *Cldn3* expression is reduced in the JEJ of female mice (**Fig.2B**, right, *p=0.008*) which are characterized by stress-induced anxiety- and depression-like behaviors^14^. Exposure to 28-d CVS is associated with the development of maladaptive behaviors in both sexes^34^ nevertheless, *Cldn3* expression was reduced only in males (**Fig.2C**, left, *p=0.03*), suggesting that other alterations might be present. Thus, like for CSDS, transcriptomic profiling was performed for genes related to tight junctions, tight junction-associated proteins, mucus layer formation and serotonin metabolism on the JEJ of unstressed controls vs male and female mice subjected to 6-d or 28-d of variable stress, revealing specific sex, stress type and duration patterns (**Fig.2D**). No difference was observed for the estrus cycle phase (**Supp. Fig.2**). In males, 10-d CSDS increased *Cldn3* only while 28-d CVS decreased expression of several tight junctions and tight junctions associated proteins. No effect was observed for mucus layer formation, serotonin metabolism or after the 6-d SCVS paradigm. In contrast, changes were mostly observed after exposure to variable stress in females (**Fig.2D**). Overall, assessment of *Cldn3* expression across stress paradigms suggested a more profound impact after 28-d CVS in males (top left, *p<0.0001;* bottom left, *p<0.0001*) vs 6-d SCVS (top right, *p=0.0071;* bottom right, *p=0.0118*) in females (**Fig.2E**). Other genes differentially impacted in a sex-specific manner by 28-d CVS included *Cldn7* (*p=0.002*), *Cldn12* (*p=0.035*) and *Ido1* (*p=0.013*) (**Fig.2F**).

**Figure 2.**
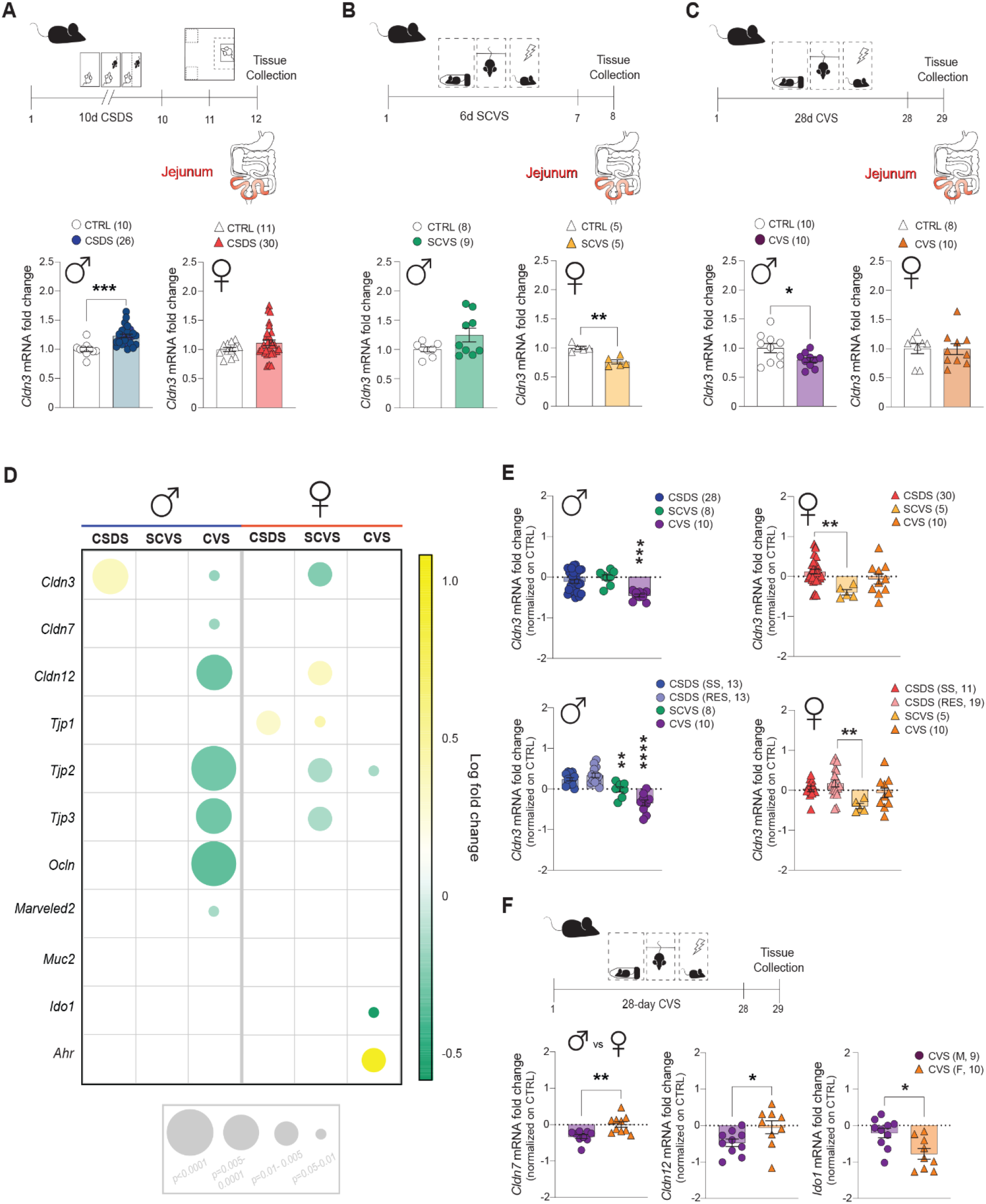
Changes in jejunum (JEJ) tight junction expression are dependent on stress type and duration. **A**) Experimental timeline of the chronic social defeat stress (CSDS) paradigm with graphs of *Cldn3* gene expression comparing control and stressed groups of male and female mice. **B**) Subchronic variable stress (SCVS) experimental timeline with comparison of *Cldn3* gene expression results below of male and female mice. **C**) Experimental timeline of chronic variable stress (CVS) with *Cldn3* gene expression results between control and stressed groups of male and female mice from this paradigm. **D**) Representation of gene expression changes in the JEJ of stressed mice across stress models in both males and females. Circle diameter represents the statistical significance of the gene expression change. Circle color represents directionality of change vs unstressed controls with green as a downregulated gene and yellow, an upregulated gene. **E**) Direct comparison of *Cldn3* gene expression changes in male and female stressed mice exposed to different stress types; CSDS, SCVS and CVS [top], with CSDS phenotypes separated to SS, RES [bottom]. **F**) Direct sex comparison of *Cldn7, Cldn12* and *Ido1* gene expression changes in mice exposed to 28-d CVS. T-tests and one-way ANOVA followed by Bonferroni’s multiple comparison test for changes between groups. \**p*<0.05, \*\**p*<0.01, \*\*\**p*<0.001, \*\*\*\**p*<0.0001.

### Detailed morphological assessments of stress-induced changes in JEJ Cldn3 expression

Next, we aimed to confirm that stress-induced alterations in tight junction gene expression are also reflected at protein level. Female mice were subjected to the 6-d SCVS paradigm then JEJ tissue was collected 24h later (**Fig.3A**). Thin 6-um slices were double stained with Cldn3 (red) and F-actin (green), and morphological analysis performed using the Imaris software (**Fig.3B-F**). Structural and organizational properties of tight junctions play a major role in their function and maintenance of the gut barrier integrity^1^. Functional tight junctions are formed when claudins interact with other transmembrane proteins, junction-associated scaffold proteins and the actin cytoskeleton^35^. Exposure to 6-d SCVS reduced Cldn3 protein level (**Fig.3C**, *p=0.0357*), in line with the changes observed at gene expression (**Fig.2B**), while no difference was measure for F-actin, a key component of the cytoskeleton (**Fig.3C**). The Imaris software has tools to evaluate colocalization of targets of interest and thus, overlap between Cldn3 and F-actin was assessed as shown on **Fig.3D**. The Pearson’s correlation coefficient revealed a significant relationship between loss in Cldn3/F-actin overlap and 6-d SCVS exposure while no change was measured with the Mander’s correlation coefficient (**Fig.3E**). Nonetheless, a trend toward lower colocalization volume was observed after stress (**Fig.3F**, *p=0.0714*).

**Figure 3.**
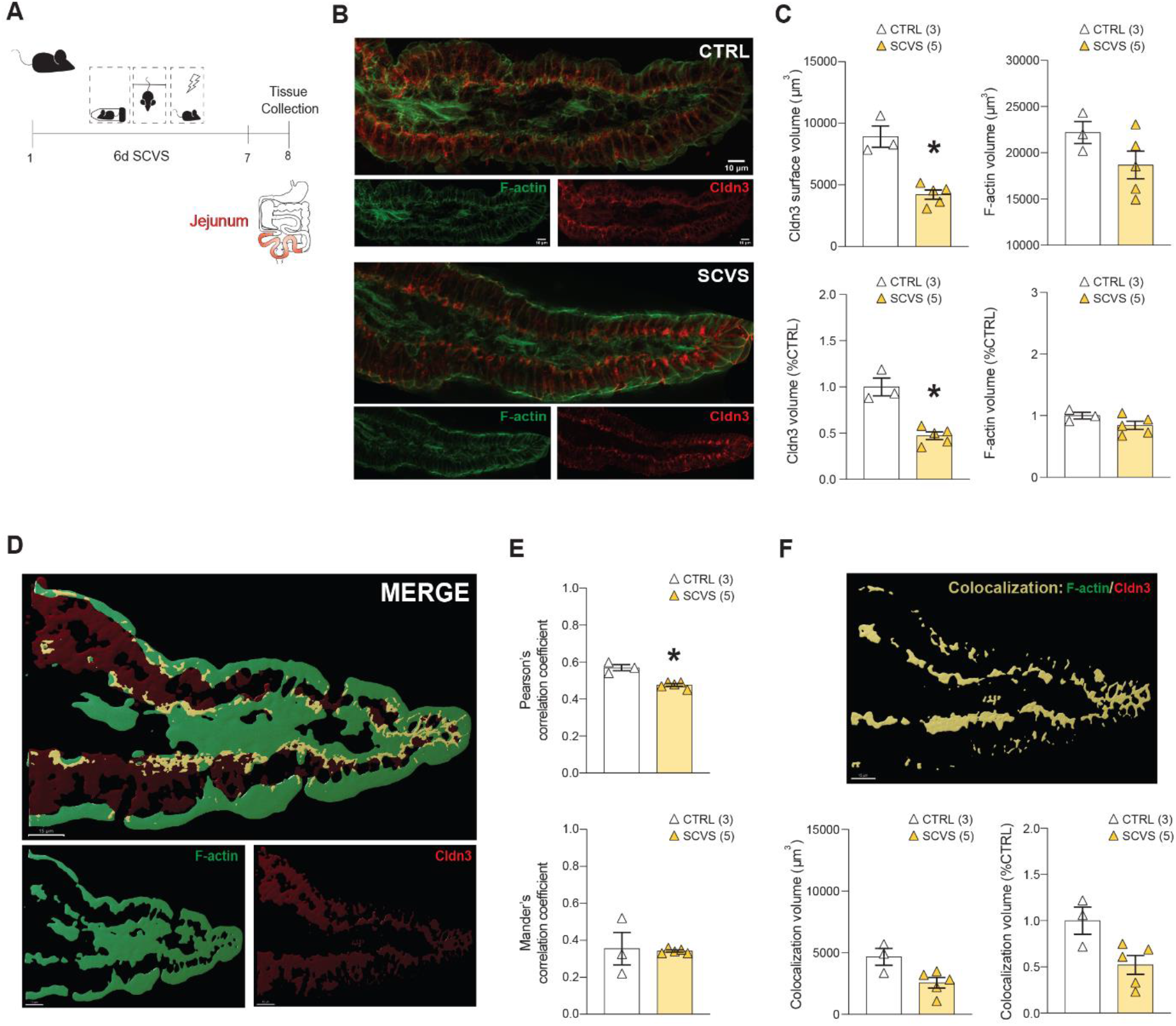
Morphological assessments of stress-induced changes in jejunum (JEJ) Cldn3 expression. **A**) Experimental timeline of female SCVS paradigm and tissue collection. **B**) Cldn3 protein level was lower in SCVS exposed mice while no difference was measured for F-actin (**C**). **D**) Representative image of Imaris volume rendering for surface volume determination. **E**) Pearson’s correlation coefficient revealed decreased colocalization of F-actin and Cldn3 signal intensities. However, no significant differences in co-occurrence of F-actin and Cldn3 was detected with Mander’s colocalization coefficient. **F**) Surface volume of colocalized regions of F-actin and Cldn3 extracted from the image in (**D**) were lower in SCVS mice without reaching significance (*p=0.0714*).

Nano-scale architecture of the BBB tight junctions has been receiving increasing attention since it could help better understand their functions and contribution in disease pathology^36^. Here, we took advantage of machine learning-based algorithms to characterize further four features of the JEJ tight junctions: ruffles, width, fragmentation, and diffusion. Images of JEJ samples from female mice exposed to 6-d SCVS (**Fig.4A**) and double stained for Cldn3 (red) and F-actin (green) were acquired (**Fig.4B**) then analyzed using a custom software. A total of 1426 image crops (controls: 452; SCVS: 974) were annotated according to the criteria defined in **Fig.4C** with a value ranging from 0 to 1. Unsupervised k-means clustering allowed to group, in a blind manner, the crops into 7 different clusters (**Fig.4D**) providing an overview and comparison of the Cldn3 tight junction properties between unstressed controls and mice subjected to 6-d SCVS (**Fig.4E**). Representative images are provided for each cluster (**Fig.4F**) along with their properties for the four feature values (**Fig.4G**). Cluster 7 (pink), which is associated with a high number of ruffles, was of particular interest as it is absent in the control mice but observed in all stressed animals (**Fig.4E**). Indeed, ruffled tight junctions are associated with increased paracellular permeability and thus, a loss of barrier integrity^35^. To our knowledge, this is the first detailed morphological characterization of the impact of stress on the gut barrier integrity. Furthermore, an increase in ruffled tight junctions suggests that stress-induced JEJ leakiness may contribute to the neuroimmune mechanisms of MDD by allowing gut-related inflammatory mediators to leak into the bloodstream.

**Figure 4.**
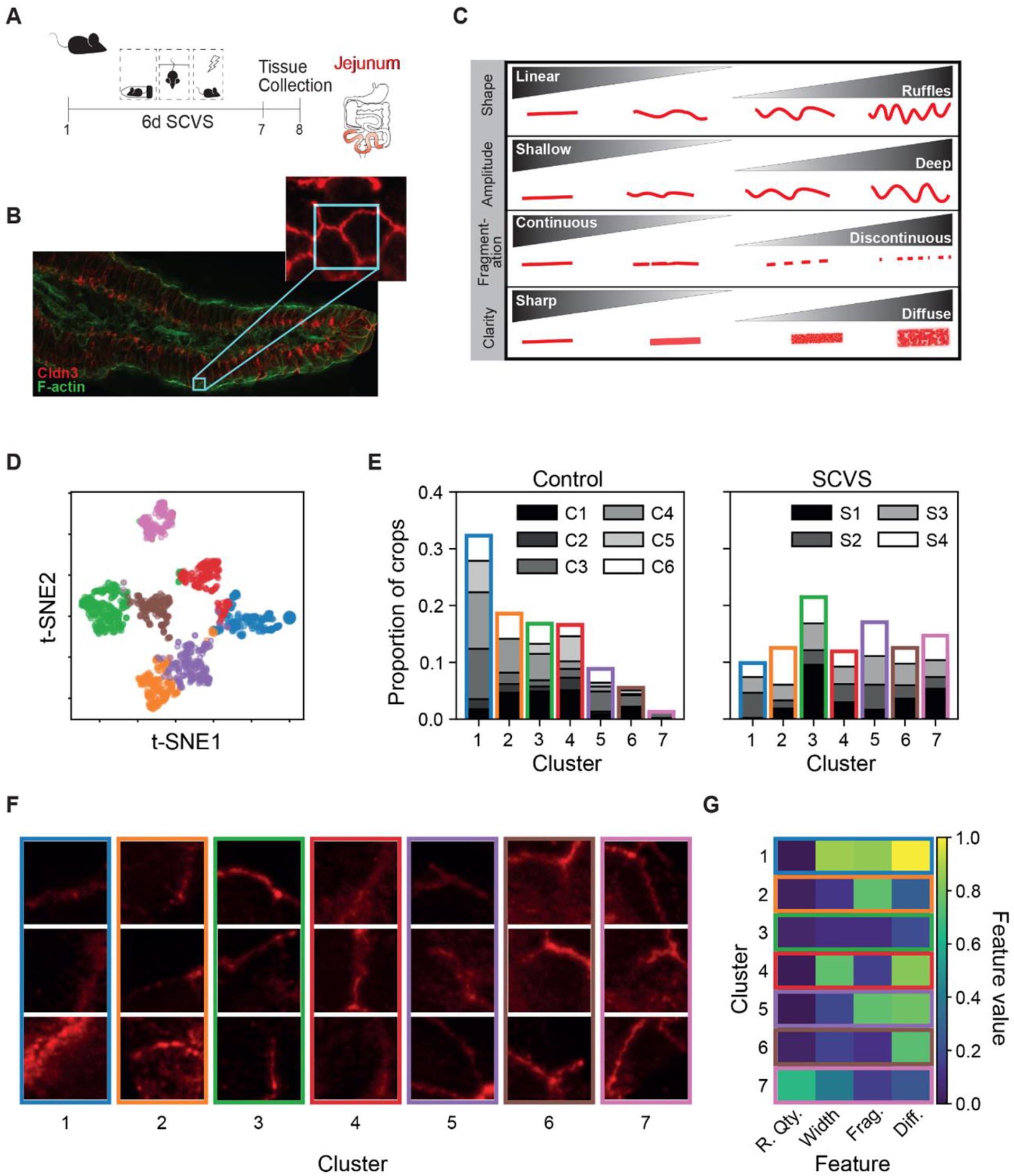
Detailed morphological assessments and k-means clustering analysis of stress-induced changes in jejunum (JEJ) Cldn3 protein expression. **A**) Experimental timeline of female SCVS paradigm. **B**) Representative immunofluorescent image of Cldn3 and F-actin. **C**) Table describing the JEJ tight junction features and parameters analyzed. **D**) t-SNE visualization of the k-means clustering. The four features (ruffle quantity, width, fragmentation, and diffusion) are projected in two dimensions using the t-SNE algorithm. Each color corresponds to a different cluster identified with k-means. Tight junction crops (5.52×5.52μm) with similar feature values are clustered together and are closer together in the t-SNE projection. **E)** Proportion of tight-junction crops from each image in each cluster for control and stressed animals. The different shades of gray correspond to different images to show that the distribution was not skewed by the overwhelming presence of a cluster in a single image. In the control condition, cluster 1 contains over half of the tight junction crops, while very few are in cluster 7. For the chronic-stress condition, there is an increase in crops belonging to clusters 5, 6 and 7. **F)** Examples of Claudin-3 tight junction crops associated with each cluster. These were selected from the 20 crops with features closest to the median point of each cluster using cosine distance. **G)** Feature “barcode” of the clusters identified with k-means clustering. Each entry corresponds to the median value of a feature in the given cluster.

### LPS-induced inflammation promotes loss of JEJ tight junctions in males only

To confirm a causal role of stress-associated inflammation in the alterations observed at the JEJ tight junctions, mice were pharmacologically treated with lipopolysaccharide (LPS). This endotoxin is a product of the outer membrane of gram-negative bacteria commonly used to study inflammation-induced behavioral changes in rodents^37^. Importantly, it has translational value since LPS is elevated in the plasma of individuals with MDD or anxiety disorders^38^, although this would need to be replicated in larger cohorts. Male or female mice received an i.p. injection of LPS (0.83 mg/kg) and after 24h samples were collected for tight junction analysis in the JEJ (**Fig.5A, D**). Treatment with LPS led to profound changes in the male JEJ gene expression with a loss of tight junctions (*Cldn3, p=0.0001*; *Cldn7, p<0.0001; Cldn12, p<0.0001; Ocln, p<0.0001; Marveld2, p<0.0001*), tight junction-associated proteins (*Tjp1, p<0.0001; Tjp2, p<0.0001; Tjp3, p=0.0001*), mucus layer-related *Muc2* (*p=0.0076*) and finally, *Ido1* (*p=0.0052*) and *Ahr* (*p=0.0016*), which are linked to serotonin metabolism (**Fig.5B-C, Supp. Fig.3B**). Conversely, no significant change was observed for female mice except for serotonin related *Ido1* (*p<0.0001*) and *Ahr* (*p=0.0115*) (**Fig.5E-F, Supp. Fig.3C-D**), suggesting that other biological mechanisms underlie stress-induced alterations in JEJ tight junctions in females.

**Figure 5.**
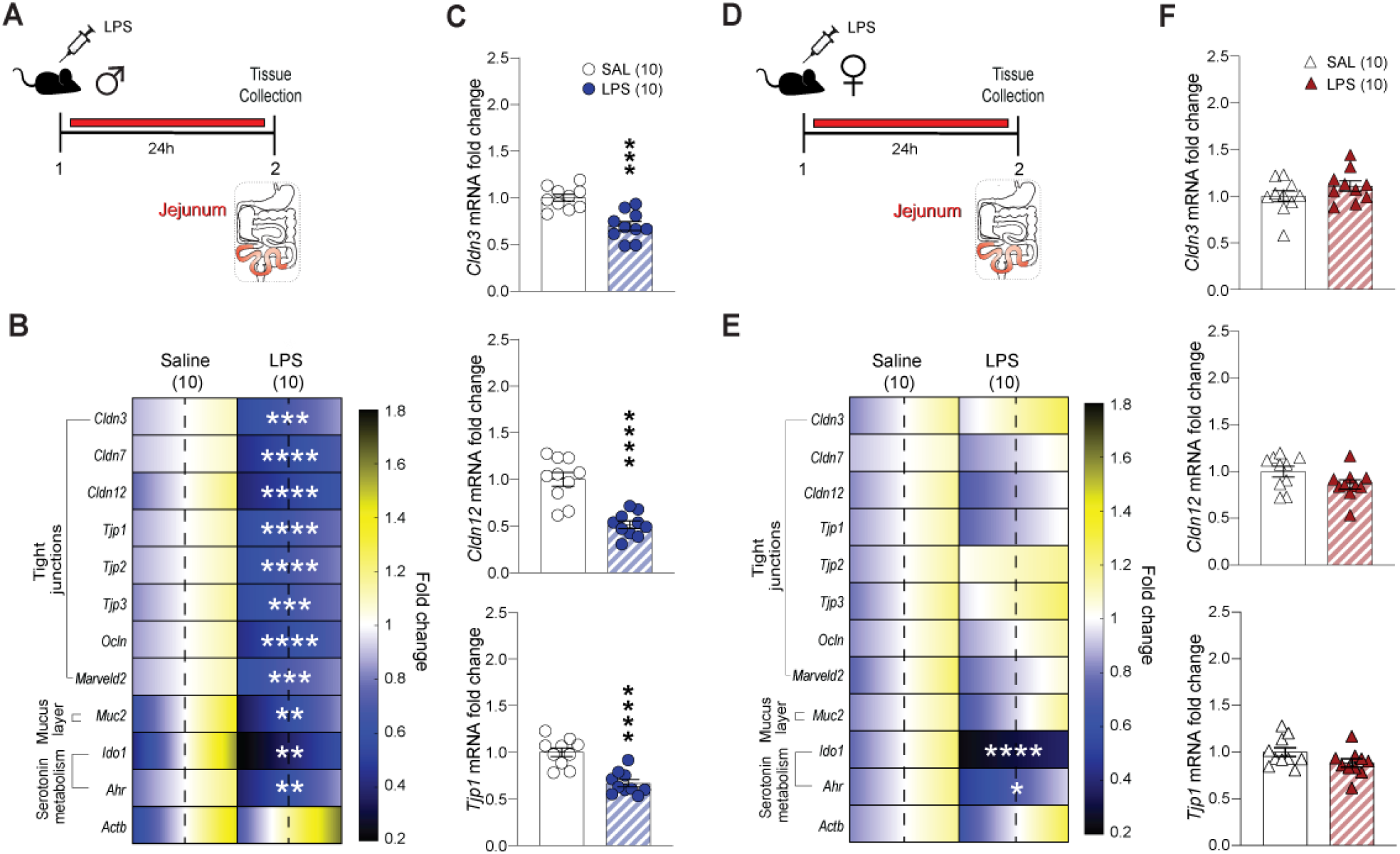
LPS-induced inflammation promotes loss of jejunum (JEJ) tight junction expression in males only. **A**) Experimental timeline of lipopolysaccharide (LPS) injection and tissue collection in males. **B**) Quantitative PCR revealed significant changes in JEJ of LPS-treated mice compared to controls (saline) for gene expression of targets related to tight junctions, the mucus layer or serotonin metabolism. The range of color indicates individual differences within a group; S.E.M. from the average represented by the dashed line. *Cldn3, Cldn7, Cldn12, Tjp1, Tjp2, Tjp3, Ocln, Marveld2, Muc2, Ido1* and *Ahr* expression was reduced in males after LPS and graphs are provided for *Cldn3, Cldn12* and *Tjp1* (**C**). **D**) Experimental timeline of LPS injection and tissue collection in females. **E**) Quantitative PCR revealed no significant changes in jejunum of LPS treated female mice compared to controls (saline injection) for gene expression of targets related to tight junctions including *Cldn3, Cldn12* and *Tjp1* (**F**). Data assessed with Mann-Whitney U test. \*\**p*<0.01, \*\*\**p*<0.001, \*\*\*\**p*<0.0001.

### Sex-specific effects of stress exposure on fecal microbiota of male and female mice

Due to the relationship between intestinal microbiota and tight junctions of the intestine^39-41^, we analyzed bacterial populations in our various stress models, for both sexes, to determine if dysbiosis may be linked to JEJ tight junction’s sex-specific changes. Male and female mice were first exposed to 28-d CVS then fecal microbiota was compared (**Fig.6A**). No difference in the alpha-diversity Shannon and Chao1 indices was observed in either sex (**Supp. Fig.4A-C**). In contrast, beta-diversity measures of dissimilarity of whole microbiota communities between groups revealed two distinct clusters for CVS male (**Fig.6B**, *p=0.042*), but not female (**Fig.6C**), mice vs unstressed controls. In line with the major changes reported for males at the JEJ tight junctions after 28-d CVS (**Fig.2)**, a loss of Bacteroidetes (*p=0.013*) and a rise of Firmicutes (*p=0.033*), the two most abundant phyla composing our fecal microbiota samples, were noted (**Fig.6D**). In contrast, no difference was noted for females exposed to 28-d CVS (**Fig.6E**), again in line with the limited number of alterations observed for tight junction expression after exposure to this stress paradigm (**Fig.2)**. CVS-exposed males also had a decrease of the *S24-7* (*p=0.016*), *Lactobacillaceae* (*p=0.011*), and *Bacteroidaceae* (*p=0.057*, data not shown) families, along with a rise in *Lachnospiraceae* (*p=0.022*) and in *Ruminococcaceae* (*p=0.039*) (**Fig.6F**).

**Figure 6.**
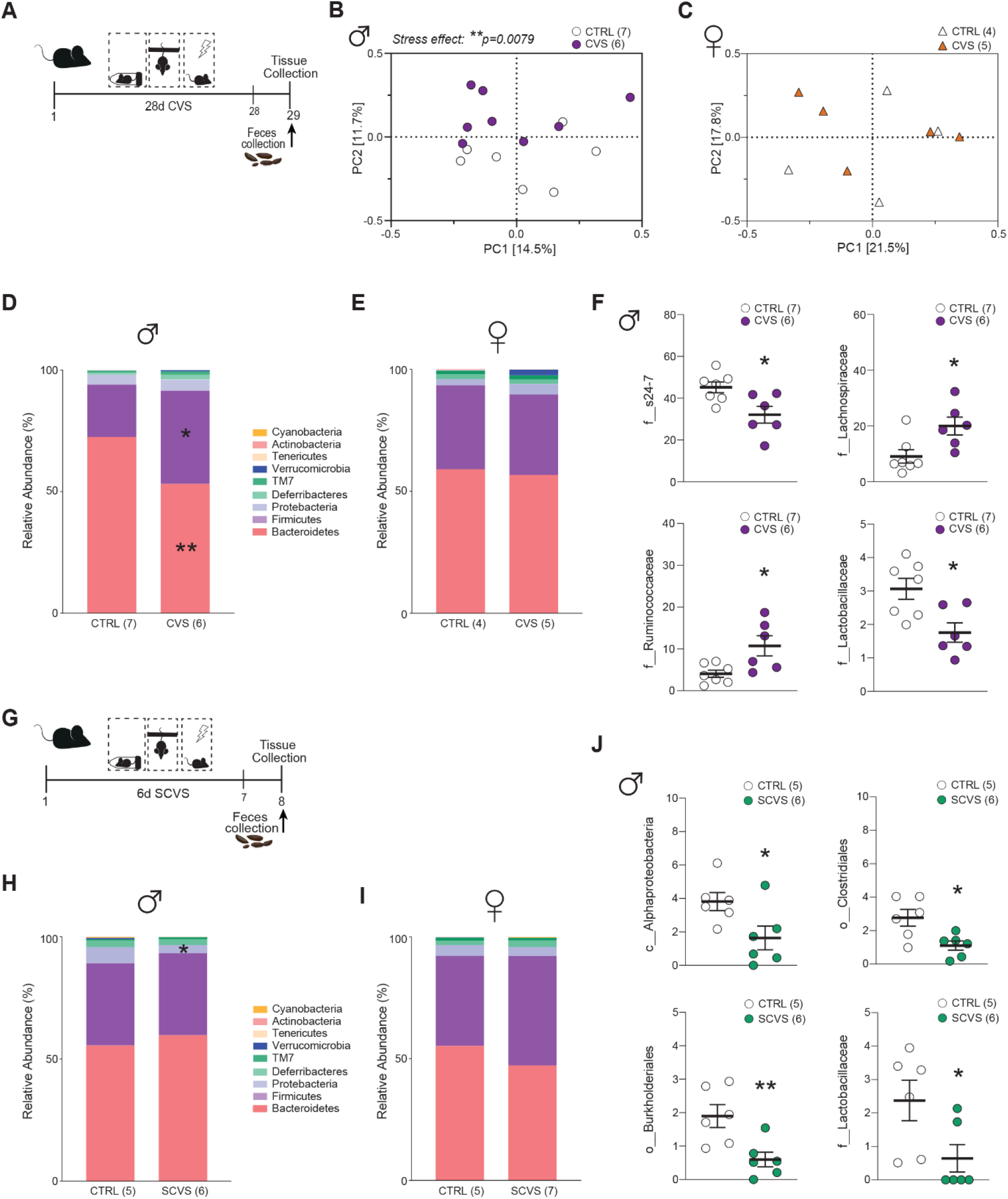
Sex-specific effects of stress exposure on fecal microbiota of male and female mice. **A**)Experimental timeline of 28-day chronic variable stress (CVS) exposure and feces collection. **B**)Analysis of beta-diversity revealed that CVS males significantly differed from unstressed controls while no difference was noted for females (**C**). **D**) Relative abundance of phylum communities showed decreased Bacteroidetes and increased Firmicutes following CVS in males with again no change for females (**E**). **F)** At the family level, CVS-exposed males had significant changes in the *s24-7, Lachospiraceae, Ruminococcaceae* and *Lactobacillaceae* families. **G**) Experimental timeline of 6-day subchronic variable stress (SCVS) exposure and feces collection. **H**) A reduction in the Proteobacteria phylum was observed in males after 6-d SCVS compared to unstressed controls. **I)** No significant change was noted for females despite exposure to the same stress paradigm. **J)** *Alphaproteobacteria* (unassigned), *Clostridiales* (unassigned), *Burkholderiales* and *Lactobacillaceae* abundances were all decreased in 6-d SCVS males. Unpaired t-tests were used for two-group comparisons. \**p*<0.05, \*\**p*<0.01.

After 6-d SCVS (**Fig.6G**), the alpha- and beta-diversity metrics did not differ between groups (**Supp. Fig.4D-H**), but the phylum Proteobacteria was decreased (**Fig.6H**, *p=0.041*) in male mice, a change that was potentially driven by reductions of the Proteobacteria members *Alphaproteobacteria* (unassigned; *p=0.035*) and *Burkholderiales* (*p*=*0.009*). SCVS-exposed males also had fewer *Clostridiales* (unassigned; *p*=*0.015*) and *Lactobacillaceae* (*p=0.026*) (**Fig.6J**). As for females, no differences were detected in relative abundances of the various phyla (**Fig.6I**), although a few changes at lower levels were apparent in this group, including an enrichment of the family *Ruminococcaceae* (*p=0.052*, data not shown) and lower levels of the genus *Alistipes* (*p=0.037*) (**Supp. Fig.4I**).

The gut microbiota has been implicated in male mice vulnerability to CSDS^42^ but, to our knowledge, it was never explored in females. Thus, female mice were subjected to 10-d CSDS^14,22^ and feces collected after stress exposure (**Supp. Fig.5A**). The Shannon and Chao1 indices and beta-diversity did not differ between groups (**Supp. Fig.5B-C**), but the examination of relative abundances throughout taxonomic ranks highlighted elevations of unassigned members of the *Bacteroidales* order following CSDS in female mice exposed to social stress (**Supp. Fig.5D-E**, *p=0.041*). Post hoc analysis revealed that SS, but not RES, mice had higher unassigned *Bacteroidales* when compared to unstressed controls (**Supp. Fig.5E**, *p=0.026*), suggesting that this subpopulation of *Bacteroidales*, although unassigned at that time, might play a role in social stress vulnerability in females.

### Blood biomarkers are associated with loss of gut barrier integrity

We recently identified blood-based vascular biomarkers, associated with inflammation of the BBB, in SS mice and women with MDD^14^. Identification of MDD-related biomarkers is greatly needed to help guide clinical diagnosis^1^. Stress-induced changes in tight junction expression and microbiota populations may reflect increased gut barrier permeability. Thus, we explored here the potential of gut-related circulating markers as indicators of stress vulnerability vs resilience. Blood was collected prior and after exposure to 10-d CSDS and the serum of male mice was analyzed for LPS-binding protein (LBP), a marker of gut leakiness^1^ (**Fig.7A, Supp. Fig.6C-E** for behavioral phenotyping). Social stress exposure induced an increase in circulating LBP in the blood serum of SS (*p=0.0033*) but not RES or unstressed control mice (**Fig.7B**, left). Moreover, LBP level was negatively correlated with social interactions (*p<0.0001*) (**Fig.7B**, right). Subtracted pre-CSDS blood LBP level was used to dampen individual differences and confirm causality with stress exposure. Next, we evaluated if this biomarker could be relevant for females as well. Blood LBP was evaluated in the serum of female mice prior vs after exposure to 6-d SCVS paradigm (**Fig.7C**). Indeed, it is in this context that we observed the most significant changes in JEJ Cldn3 tight junctions (**Fig.2D**). Like in males, stress exposure was associated with elevated circulating LBP when compared to unstressed controls (*p=0.0188*) (**Fig.7D**). Baseline LBP was not different between unstressed control males and females (**Fig.7E**). Finally, we evaluated translational value of LBP as a potential biomarker of mood disorder by measuring it in blood serum samples from individuals with MDD. High circulating LBP was observed for men and women with MDD vs controls, without reaching significance (**Fig.7F**). However, consideration of sex revealed an increase in women (*p=0.0434*), but not men, with MDD (**Fig.7G**). Like in mice, no difference was noted for samples of the men and women in the control group (**Fig.7H**). These findings suggest an increase in gut barrier permeability following stress exposure in mice and possibly, in individuals with MDD.

**Figure 7.**
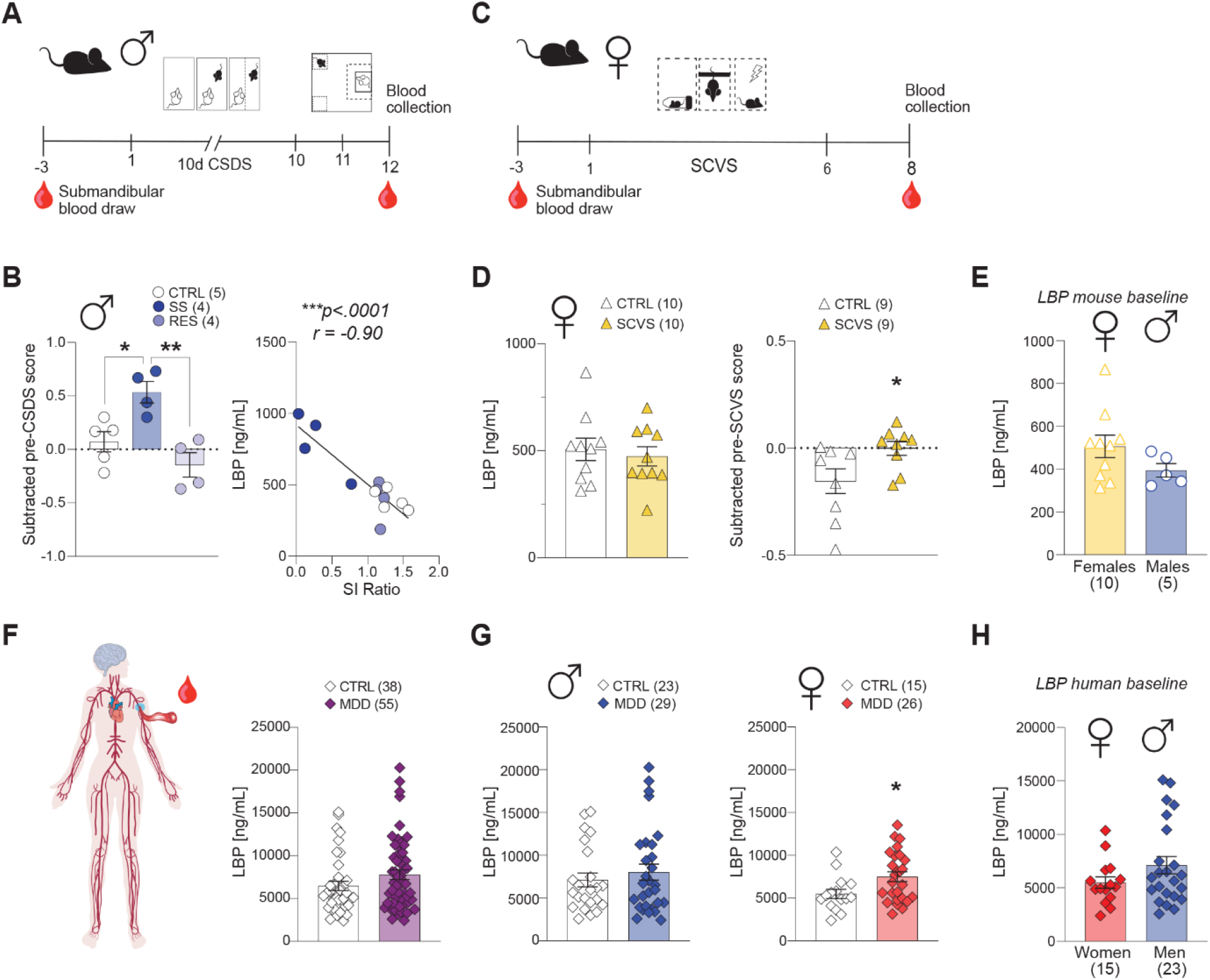
Blood biomarkers are associated with loss of gut barrier integrity in stressed mice and individuals with major depressive disorder (MDD). **A**) Experimental timeline of 10-day chronic social defeat stress (CSDS) and blood collection prior and after stress exposure. **B)** LPS-binding protein (LBP) is increased in stress-susceptible (SS), but not resilient (RES) male mice when compared to unstressed controls (CTRL) after CSDS and negatively correlated with social interaction (SI) ratio. **C)** Experimental timeline of 6-d subchronic variable stress (SCVS) and blood collection prior and after stress exposure. **D)** Circulating LBP appears similar between unstressed and stressed groups of female mice but is in fact increased after 6-d SCVS when LBP level is compared after vs prior stress. **E)** Baseline blood LBP is higher in unstressed control female mice when compared to males of the same group without reaching significance. **F)** Circulating LBP is upregulated in individuals with MDD, an effect driven by women (**G**). **H)** Baseline blood LBP in healthy control women and men is similar. T-tests and one-way ANOVA followed by Bonferroni’s multiple comparison test for changes between groups. Human data were assessed with two-tailed Mann-Whitney U test. \**p*<0.05, \*\**p*<0.01, \*\*\**p*<0.001.

## Discussion

A growing body of literature connects aberrant gut-brain axis signaling in MDD, nevertheless, a direct link between gut permeability and BBB leakiness, facilitating passage of circulating inflammatory mediators into the brain in this context is still debated^1^. The BBB and gut barrier had not been compared directly until a recent evaluation of female rats facing social isolation stress, which demonstrated shared modifications to *Ocln, Tjp1* and *Cldn5* gene expression in the prefrontal cortex and the ileum^43^. Considering our recent work identifying sex-specific BBB tight junction changes in response to social defeat^12,14^ and variable stress^14^, we examined how these models influence gut integrity. Overall, our results demonstrate altered expression of JEJ tight junctions following stress, dependent on the type and duration of exposure. Indeed, alterations had patterned sex-specific changes, consistent with the knowledge that female and male mice are disparate in their behavioral and biological response to various stress models^25,44^. LPS treatment decreased many tight junction proteins in the intestine of males exclusively, highlighting the potential role of inflammatory pathways on the gut. Measuring serum marker LBP provided indirect evidence of increased intestinal permeability in both sexes. Microbiota sequencing revealed sex-specific altered microbial populations post-stress which may be linked to the manifestations of different changes in intestinal permeability along with depressive-like behaviors.

Recurrent alterations in Cldn3 expression, part of the “tightening” group of tight junction proteins^33,45^, suggests that chronic stress exposure affected barrier permeability. In rodents, Cldn3 expression relates to functional measurements of permeability^46^ and *in vitro* overexpression reduced paracellular ion and large-molecule permeability^47^. Cldn3 alterations are implicated in inflammatory bowel and celiac diseases^48^, though precise functions and roles in pathology are unclear. Our findings that CSDS elevated *Cldn3* expression in males aligns with previous reports that intestinal Cldn3 promotion is a repair initiative by the host in response to epithelial barrier injury^49^. On the contrary, 28-day CVS caused a *Cldn3* reduction, which implies the reparative strategy is not sufficient to overcome a longer-term stress exposure.

Imaging showed Cldn3 protein decrease in females reflecting mRNA expression changes. Shifts in claudin expression are not simply restricted to protein or RNA quantity, as tight junction structures and functions are highly dynamic. Tight junction borders typically display as linear structures between cells, however, adjustments in junctional assembly and interactions with scaffolding and cytoskeletal proteins alter these morphologies^35,50^. Briefly, ruffles present as a zig-zag shaped border and may be indicative of enhanced paracellular permeability^35^. Increased ruffling of tight junction borders following SCVS supports our previous statement that alterations in Cldn3 expression is likely indicative of disrupted barrier function in these animals.

The relationship between inflammatory bowel diseases and MDD has promoted clinical investigations into biomarkers of intestinal dysfunction in the context of depression^30,31,38,51^. However, limited investigations exist in the context of chronic stress models. Here, elevated serum LBP predominated in SS, and not RES, males following CSDS, highlighting its potential as a sex-specific biomarker for vulnerability to this form of stress. LBP rises provides indirect evidence of microbial translocation, due to its specificity to LPS endotoxin. To our knowledge, this is the first report of this intestinal-related biomarker in chronic stress models. In accordance with previous clinical reports^30^, the observed serum LBP levels were elevated in MDD patients. However, this effect was specific to women only in our cohort, and we did not find other reports that considered MDD and compared the effect between sexes.

As for MDD, women have a higher risk of developing inflammatory bowel diseases^1^, therefore, tight junction alterations were expected in females following LPS-induced inflammatory challenge. Surprisingly, tight junction gene expression was unaltered in females, while considerable decrease occurred in male mice. Certainly, many gene markers implicated in gastrointestinal disorders or claudins with still undetermined functions could be changing in a sex-specific manner. Novel roles for peripheral serotonin metabolism are implicated in inflammatory, immune, and metabolic signaling pathways^52^, prompting the inclusion of serotonin pathway molecules *Ido-1* and *Ahr* to the intestinal gene expression analysis. LPS treatment reduced levels of both serotonin-related molecules in females without affecting tight junctions like for males, highlighting a potentially female-specific pathway. Indeed, progesterone modulates the colonic serotonin system and in women with inflammatory bowel diseases, both the serotonin transporter and serotonin levels are diminished^53^. IDO-1 is highly upregulated in the human gut epithelium during inflammation^54^ and linked to symptoms in MDD^55,56^. However, the reduction following LPS implies another mechanism at play. AHR responds to microbial metabolites in an attempt by the host to resist colonization by opportunistic pathogens^57^, therefore, these changes may reflect microbiota population changes.

LPS can interact with tight junctions directly^41^ with implications for both the gut barrier and BBB. Indeed, in mice, the absence of gut microbes is associated with increased permeability of the BBB^58^. Furthermore, traumatic stress models induce shifts in microbiota diversity and intestinal inflammation with downstream effects on hippocampal Cldn5^59^. Throughout the examined stress paradigms, only the CVS-exposed males differed from unstressed controls in terms of beta-diversity. This group also had microbiota shifts at the phylum level similar to those seen in individuals with irritable bowel syndrome^60^, for whom Cldn3 and Ocln reductions have also been reported^61^. Compositional changes in CVS males also coincides with chronic unpredictable mild stress exposure in rats, especially those changes within the family *Lachnospiraceae*^*62*^. Although *Lachnospiraceae*, along with *Lactobacillaceae* and *Ruminococcaceae*, have the ability to produce butyrate and other short-chain fatty acids able to strengthen the intestinal barrier through up-regulation of tight junctions^63,64^, a number of genera of *Lachnospiraceae* have been implicated in intra- and extra-intestinal diseases^63^ and thus future studies will be necessary to decipher the mechanisms involved. As for short-term CVS exposure, decreases in taxa with pro- as well as anti-inflammatory properties in the intestinal environment (e.g., decreases in both the pro-inflammatory Proteobacteria and the anti-inflammatory *Lactobacillaceae*) were apparent in male mice. Proteobacteria elevations have been reported in male mice exposed to longer or more severe stress^42^, and thus the reductions in 6-day SCVS males could represent a temporary stress-induced compensatory change. Intriguingly, enteral administration of *Lactobacillus* probiotic species can modulate intestinal Cldn3 expression^49^.

Limitations of this study include investigating only the JEJ and feces instead of gut content, which eliminates the potential of region-specific identifications. Reports of stress effects on jejunal permeability are scarce, making comparisons within the literature a challenge. Most focus on the colon due to inflammatory bowel disease pathogenesis, however, the concept of intestinal permeability in MDD has not been associated with a single region of the gastrointestinal tract. The functional roles of the jejunum and the relevancy of small intestinal dysfunctions highlight the value of further investigations^27,28^. Another potential drawback involves performing the CSDS paradigm, in which physical injuries are a concern for potential stimulation of the immune and inflammatory system. Physical examinations of animals were performed at time of sacrifice to observe the number of wounds to consider this as a potential confounder including for peripheral measurements of LBP. However, previously repeated social disruption stress and restraint stress induced bacterial translocation to the mesenteric lymph nodes independently of wounding condition^65^.

To sum up, our study shows that chronic social or variable stress can induce gut barrier alterations in male and female mice. Changes in JEJ tight junctions along with microbiota populations are sex-, type- and duration-dependent, which might be associated to the differences reported in MDD symptomatology. To our knowledge, we provide the first detailed morphological characterization of the JEJ tight junctions following stress exposure by applying novel tools and algorithms that will be freely available and could be applied to various conditions including inflammatory bowel diseases. Finally, by focusing on the small intestine this project brings novel insights into the biology underlying stress responses in mice and inform on potential biomarkers of MDD such as circulating products related to gut barrier leakiness.

## Methods

### Animals

Experimental mice were naïve male (∼25g) and female (∼20g) C57BL/6 mice, 7–9 weeks of age at arrival (Charles River Laboratories, Québec, Canada). Sexually experienced retired male CD-1 breeders (∼40 g), 9-12 months of age were used as aggressors (AGG), residents for the social defeat procedures (Charles River Laboratories, Québec, Canada). All mice were group housed in 27 × 21 × 14 cm polypropylene cages upon their arrival and left undisturbed for one week of acclimation at the housing facility of CERVO Brain Research Center prior to any procedures commencing. Mice maintained on a 12-h light–dark cycle (lights on from 0800 to 2000 h) with temperature (22 °C) and humidity (63%) kept constant were provided free access to water and food (Teklad Irradiated Laboratory Animal Diet, Madison, USA). All experimental procedures were approved by the animal care and use committee of Université Laval (2018-052) and met the guidelines set out by the Canadian Council on Animal Care.

### Chronic Social Defeat Stress (CSDS)

As previously described^21^, in the CSDS model, a C57BL/6 mouse is repeatedly subordinated by an AGG mouse for daily bouts of social stress. Before the experiment, AGG mice are screened for aggressive behaviours against a C57BL/6 mouse for 3 days. Then the AGG is designated a social defeat cage (26.7 cm width × 48.3 cm depth × 15.2 cm height, Allentown Inc.) separated in half with a clear perforated Plexiglas divider (0.6 cm × 45.7 cm × 15.2 cm). The experimental C57BL/6 mice are placed in the home cage of an unfamiliar CD-1 male for bouts of physical stress lasting 5 minutes daily over a period of 10 consecutive days. After each physical stress period, the mouse is returned to the other side of the clear plastic divider, leaving the experimental mouse exposed to overnight sensory contact with the AGG mouse. Control animals were housed 2 per social defeat cage, one each side of the Plexiglas divider and kept in the same room as experimental mice. After the last day of social defeat, the experimental mice are single housed for 24 hours before conducting the social interaction (SI) test.

In the female CSDS paradigm, the procedure is adjusted as described by Harris *et al*.^22^. Before each defeat, ∼60ul of urine collected from a CD-1 male was applied to the tail base and vaginal region of the female mouse. The odor from the urine induces dominant behaviour from the AGG mouse. For urine collection, CD-1 mice were placed in metabolic cages (Life Science Equipment) during the dark phase of the light/dark cycle. Urine was collected the following morning, filtered, aliquoted in 0.5 mL tubes and stored at −80°C until use.

### Social Interaction (SI) Test

Following CSDS, mice are characterized for vulnerability to stress by way of SI test to establish behavioural phenotypes^21^. In this test, the mouse’s propensity to socialize is evaluated. In two trials, this test assesses exploratory behavior of mice in an open field, first alone and then in the presence of an unfamiliar CD-1 mouse contained in a small wire cage within the open field (42 cm x 42 cm x 42 cm, Nationwide Plastics). Movements are tracked by an automated system (AnyMaze™ 6.1, Stoelting Co.) during each 2.5-minute trial. The time spent in the interaction zone, the region surrounding the small cage, compared to the rest of the arena is assessed. The SI ratio is the score obtained by dividing the time spent in the interaction zone with AGG present divided by the time when AGG is absent. Equal time spent when in Trial 1 and Trial 2, giving an SI ratio of 1, is typically used as the cutoff point, dividing those mice with a ratio below 1.0 to be classified as stress-susceptible (SS), while mice with a ratio above 1.0 are considered resilient (RES)^21^.

### Chronic Variable Stress (CVS)

As previously described^34^, mice are exposed to a series of alternating variable stressors (restraint stress, tail suspension, foot shocks) for 1 hour per day unpredictably for 28 days. Stressors were administered as follows: 100 mild foot shocks of 0.45mA at random intervals for 1 h (10 mice/ chamber), a tail suspension stress for 1h and restraint stress, where animals are placed inside a 50ml falcon tube, for 1h within the home cage. After day three the stressors restart with foot shocks and cycles through in this pattern for 28 days.

### Subchronic Variable Stress (SCVS)

The SCVS paradigm consists of the same procedure as CVS but for a total duration of 6 days only^14,25^. Male and female C57BL/6 mice were used as experimental mice and stressors were administered as previously described for the CVS paradigm. Stressors were administered daily for 6 days, as follows: 100 mild foot shocks for 1h (days 1 and 4), a tail suspension stress for 1h (days 2 and 5) and restraint stress for 1h (days 3 and 6).

### Tissue collection

Blood samples were collected 72h before the start of the stress protocol. Blood was collected by the submandibular bleeding method and left at room temperature in nuclease-free microtubes for 1h before processing for serum extraction. Samples were centrifuged for 2 min at 10 000 RPM at which point the separated serum was collected and transferred into a new tube. This process is repeated with centrifugation at 10 min at 3000 RPM and supernatant was collected. Serum was stored at −80°C for subsequent determination of protein levels.

Blood, feces, and tissues from the same mice were collected 24h (for SCVS or CVS) or 48h (for CSDS) after the last stressor. Trunk blood was collected after euthanasia by rapid decapitation. 1-2 fecal pellets were collected from each mouse by voluntary defecation into a sterile microtube (Eppendorf, Germany) and fecal samples were immediately placed on dry ice. Post-sacrifice, the small intestine is removed, placed in a petri dish on ice and cut into 3 sections. The most anterior portion and distal section is discarded, and the mid-section (jejunum) is kept and flushed with 0.1 M phosphate-buffered saline (PBS 1X). Two biopsies were taken with Unicore 2.00mm punch (Harris, 7093508) from intact intestinal segments, placed immediately in nuclease-free microtubes on dry ice and stored at −80 °C for subsequent analysis. The remaining intestinal segment is prepared for subsequent protein analysis following a modified version of the swiss roll protocol^66^. Briefly, intestinal segments were cut open longitudinally along the mesenteric line. Tissue is rolled over a wooden skewer with mucosa facing outwards and then transferred into a tissue mold filled with Tissue-Tek® O.C.T. Compound (Sakura, NC1862249) to embed samples. The tissue is snap frozen in isopentane on dry ice and stored at −80°C.

### Transcriptional profiling of mouse tissue

Quantitative polymerase chain reaction (qPCR) using SYBR green chemistry was performed on JEJ tissue samples for evaluation of gene expression changes of targets related to intestinal permeability, such as tight junctions, tight junction associated proteins and inflammatory markers. Total RNA was extracted with TRIzol (Invitrogen, 15596026) homogenization and chloroform layer separation. Tissues were processed using PureLink® RNA Mini Kit (Invitrogen, 12183018A) following manufacturer’s protocol for Purifying RNA from Animal Tissues. Yields and purity (ratio of absorbance at 260 and 280 nm) of extracted RNA was assessed by NanoDrop 2000 spectrophotometer (Thermo Fisher Scientific, ND-2000). Complementary DNA (cDNA) was synthesized using Maxima™ H Minus cDNA Synthesis Master Mix, with dsDNase (Thermo Fisher Scientific, M1681) from 5μg of RNA. The cDNA was applied as a template for qPCR reaction using PowerUp SYBR Green Master Mix (Applied Biosystems, A25742) containing ROX™ Passive Reference Dye. QPCR was performed with Applied Biosystems QuantStudio 5 Real-Time PCR System (ThermoFisher Scientific, MA, USA). Oligonucleotide primers are listed in **Table S1** and primers that amplify *Actb* and *Gapdh* were used as reference genes. Analysis was done by way of the ΔΔCt method.

### Immunohistochemistry of Claudin-3 (Cldn3)

Swiss rolls of JEJ tissue samples from male CSDS mice were sectioned on a cryostat (CryoStar™ NX50 cryostat, Thermo Scientific), cut into 7 μm thick sections at −17°C and mounted on Superfrost Plus slides. Slices were rinsed in PBS 1X and incubated for 30 mins in blocking solution, consisting of 10% normal donkey serum in PBS 1X. Slides were incubated overnight at 4°C with primary antibodies (**Table S2**) in solution, 1% bovine serum albumin (Life Sciences, SH3057401) and 0.01% Tween 20 (Fisher BioReagents, BP337-100) in PBS 1X. The slices from male mice were double stained with primary antibody CD326 for visualization of epithelial cells and tight junction Cldn3. Sections were washed three times with PBS 1X and incubated with secondary antibodies (see **Table S2**) for 1h at room temperature. Slices were washed three times in PBS 1X and stained with 4′,6-diamidino-2-phenylindole (DAPI) for nuclei visualization. Finally, slides were mounted with ProLong Diamond Antifade Mountant (Invitrogen, P36961), and cover slipped. Six 0.5μm thick *z*-stack images were acquired using an Axio Observer.M2 microscope (Carl Zeiss) with a 20X objective. Processing of images was done with Imaris version 9.7.2 (Bitplane, Zurich, Switzerland) for volume quantification and intensity colocalization.

Two JEJ tissue swiss roll slices per female mouse were double stained with primary antibodies for Cldn3 and actin filaments (F-actin) with the same protocol as described above. Ten 0.250 μm thick z-stack by six tiles were acquired with a 40X lens. Images were imported into Imaris for 3D reconstruction. With the Surface tool based on the F-actin staining, a reconstruction of the intestinal epithelium was created as a region of interest. A surface was also created based on the Cldn3 channel to detect the surface volume within the region of interest as well as the intensity of Cldn3 expression. Masks were made of the volume renderings and colocalization analysis was performed.

### Machine-learning based morphological analysis

A total of 10 images acquired as described in the Immunohistochemistry of Claudin-3 were analyzed, with 6 from control animals and 4 from SCVS animals. Each of these images were cropped to contain one villus, averaging a field of view of around 50 000 μm^2^. Images were annotated using a custom software (https://github.com/FLClab/junction_annotator). For the annotations, 5.52 × 5.52μm (64 × 64 pixels) image crops (hereafter: crops) were extracted from regions of interest corresponding to the external border of the villi. The border of the villi was identified using an intensity-based foreground mask from which an erosion of 300 pixels was subtracted. Pixels were assigned to the foreground if their value was above 0.75 times the mean intensity of the image. Crops were selected for annotation if at least 10% of their pixels were part of the border. A total of 1426 crops (control: 452 crops, SCVS: 974 crops) were identified by an expert to contain a structure belonging to a tight junction and were used in the analysis.

The expert annotated each image crop based on four qualitative features: (1) Ruffles Quantity, (2) Width, (3) Fragmentation and (4) Cldn3 Diffusion (**Fig.4C**). Each of these features were given a continuous value ranging from 0 to 1 by the expert using the homemade annotation program. Ruffles Quantity or Shape (1) refers to the number of ruffles in the crops belonging to the tight junction, where a value of 0 means no ruffles are visible while a value of 1 is the maximal amount of ruffling in the samples. Width or Amplitude (2) refers to the width of the Cldn3 signal where a value of 0 would be associated with a thin (diffraction-limited) line. If ruffles are visible, this value refers to the amplitude of the ruffles. Fragmentation (3) described the presence of discontinuous fragments in the Cldn3 fluorescence signal, where a value of 0 means a perfectly continuous junction. Diffusion or Clarity (4) refers to the apparent diffusion of the Cldn3 molecules at the tight junction, where a value of 0 means that the junction appears very sharp, while a value of 1 means the junction appears diffuse, or blurry.

Unsupervised k-means clustering algorithm^67^ was used to group the crops into clusters using the described 4-dimensional feature space, with clusters determined using silhouette score analysis^68^. A 2D projection of the feature space was created using the t-SNE algorithm^69^ for an easier visualization of the clustering. The proportion of crops within each cluster was calculated for control and SCVS samples to reveal potential differences in tight junction populations between both conditions (https://github.com/FLClab/junction_analysis).

### Microbiota analysis

Fecal DNA was extracted using a Stool DNA Isolation Kit (Norgen Biotek, 27600) following manufacturer’s protocols. Extracted DNA yields and purity were assessed using a Qubit Fluorometer (Invitrogen, Q33238). The V3 and V4 hypervariable regions of the 16S ribosomal RNA (16S rRNA) gene were amplified using the primers S-D-Bact-0341-b-S-17 (F: 5’ TCG TCG GCA GCG TCA GAT GTG TAT AAG AGA CAG CCT ACG GGN GGC WGC AG) and S-D-Bact-0785-a-A-21 (R: 5’ GTC TCG TGG GCT CGG AGA TGT GTA TAA GAG ACA GGA CTA CHV GGG TAT CTA ATC C)^70^. After tagging the resulting amplicons with Illumina nucleotide sequencing adapters and dual-index barcodes, the pooled library was sequenced using a 600-cycle MiSeq Reagent Kit v3 and a MiSeq Illumina system a as per manufacturer’s instructions (Illumina, San Diego, CA, USA). The resulting data was processed using QIIME 2^71^, with a median quality score of Q>30, and further analyzed using DADA2^72^. MicrobiomeAnalyst was used to calculate diversity indices from the pre-processed data using methods as previously described^73^. The parameters evaluated were the Chao1 and Shannon alpha-diversity indices and the Bray-Curtis dissimilarity beta-diversity index, the latter being followed by a Permutational Multivariate Analysis of Variance (PERMANOVA) and visualized using Principal Coordinate Analysis (PCoA). The relative abundance of bacteria at various taxonomic levels was calculated for each sample after aligning reads to taxa using the Greengenes database^74^.

### LPS treatment

Male and female mice received an intraperitoneal (i.p) injection of LPS (LPS from E.coli, O127:B8, Sigma) at 0.83 mg/kg. LPS was dissolved in sterile normal saline solution (0.9 %) and administered in a volume of 10 ml/kg of the body weight of the mice. LPS or saline (vehicle group) was administered at time zero and after 24h samples were collected for tight junction analysis in the jejunum.

### Murine ELISA and Multiplex assays of gut leakiness

The quantitative detection of serum lipopolysaccharide binding protein (LBP) was performed by Mouse LBP Enzyme-Linked Immunosorbent Assay (ELISA) kit (Abcam, ab213876) according to the manufacturer protocol. The serum was diluted 1:100 for LBP detection and the optical density (OD) of the plate was read at 450 nanometers (nm) by an Eon Microplate Spectrophotometer (BioTek Instruments Inc., Winooski VT). Data was calculated from a serial dilution curve using Gen5 Data Analysis Software and samples with a coefficient of variation above 15% were eliminated from the analysis.

### Human serum sample collection

All human blood samples were provided by Signature Bank from the Centre de recherche de l’Institut universitaire en santé mentale de Montréal (CR-IUSMM) under approval of the institution’s Ethics Committee. Samples were collected at the emergency room of the Institut universitaire en santé mentale de Montréal of CIUSSS de l’Est-de-Montreal from depressed volunteers and at the CR-IUSMM from healthy volunteers. All subjects were evaluated for depressive behaviors by the Patient Health Questionnaire (PHQ-9), which scores each of the nine Diagnostic and Statistical Manual of Mental Disorders (DSM) IV criteria^75^. Exclusion criteria involved subjects with known history of drug abuse. The demographic characteristics associated with each sample are provided in **Supplementary Table 3**. All experiments were performed under the approval of Université Laval and CERVO Brain Research Center Ethics Committee (#2021-2200).

### Human serum ELISA of gut leakiness marker

Human serum levels of LBP were assayed using the Human LBP ELISA kit standard sandwich enzyme-linked immunosorbent technology and following the manufacturer’s protocol (Abcam, ab213805). Briefly, the sample diluent buffer was used to dilute standards and samples (1:1000) before applying to each well and incubating for 90 mins. After the plate contents were discarded, anti-Human LBP antibody was added to each well before the second incubation for 60 mins. Three washes were completed with PBS 1X and the complex solution was added for a 30 min incubation. The plate was washed five times and the Horseradish peroxidase (HRP) substrate solution was added prior to the final incubation of 30 mins in the dark. All the incubation periods occurred at 37°C. Finally, the Stop Solution was added, and the OD of the plate was read at 450 nm by the Eon Microplate Spectrophotometer. Again, data was calculated from a serial dilution curve using Gen5 Data Analysis Software and samples with a coefficient of variation above 15% were removed.

### Statistical Analysis

All data are presented as means ± standard error of the mean (S.E.M.). Comparisons between group was performed using t-test, one-way ANOVAs and two-way ANOVAs with Bonferroni post hoc follow up test when required. Values of *P<.05* were regarded as statistically significant. Graphs and statistics were generated using GraphPad Prism Software version 8 (GraphPad Software Inc.). Normality was determined Shapiro–Wilk tests and Levene’s test for homogeneity of variances. Data was analyzed using the nonparametric Kruskal-Wallis test when it did not meet the assumption of normality, or the Mann-Whitney U test for pairwise comparisons. Visual representation of average and S.E.M. with heatmaps was created using Matlab-based software. Individual values were used to compute correlation matrices and p-values were determined by Matlab-based software (MathWorks). Comparison of microbiota alpha diversity was performed with Microbiomeanalyst using the phyloseq package^76^ and further comparisons using the Mann-Whitney U test. Beta diversity was performed using Bray–Curtis (dis)similarity matrices using the phyloseq package. The principal coordinate analysis (PCoA) was performed to visualize the distance matrix and the importance of the changes at the community level was assessed using permutational multivariate analyses of variance (PERMANOVA) tests.

## Acknowledgements

This work was supported by the New Frontiers in Research Fund (Exploration Grant to C.M. and A.D.), Sentinel North Initiative funded by Canada First Research Excellence Fund (Research Chair on the Neurobiology of Stress and Resilience to C.M., 2020-2024 Major Call for Proposal to F.L.C. and C.M., Postdoctoral fellowship to F.N.K.), Fonds de recherche du Quebec (FRQS) – Health (PhD scholarships to L.D.A. and K.A.D., junior 1 and junior 2 salary awards to C.M.), the Natural Sciences and Engineering Research Council (Discovery Grant to M.C.A.), and the Canadian Institutes for Health Research (PhD scholarship to L.D.A., MSc scholarship to N.O. and S.P., Vanier PhD scholarship to J.K.S., Project Grant to C.M.).

## Author Contributions

E.D. and C.M. designed research; E.D., L.D.A, F.C.R., S.E.J.P., F.N.K., J.L.S., R.G., K.A.D. and M.L. performed research including behavioural experiments, pharmacological treatments, molecular, biochemical, and morphological analysis; R.B., R.O.A., A.D., F.L.C. developed the detailed morphological analysis algorithms and pipeline; fecal sample collection and microbiota sequencing and analysis were performed by E.D. with N.O., J.K.S. and supervised by M.C.A.; the Signature Consortium contributed the human blood samples and related demographic data; E.D. and C.M. analyzed the data and wrote the manuscript which was edited by all authors.

## Signature Consortium – list of members

Frederic Aardema^5^, Lahcen Ait Bentaleb^5^, Janique Beauchamp^5^, Hicham Bendahmane^5^, Elise Benoit^5^, Lise Bergeron^5^, Armando Bertone^5^, Natalie Bertrand^5^, Felix-Antoine Berube^5^, Pierre Blanchet^5^, Janick Boissonneault^5^, Christine J. Bolduc^5^, Jean-Pierre Bonin^5^, Francois Borgeat^5^, Richard Boyer^5^, Chantale Breault^5^, Jean-Jacques Breton^5^, Catherine Briand^5^, Jacques Brodeur^5^, Krystele Brule^5^, Lyne Brunet^5^, Sylvie Carriere^5^, Carine Chartrand^5^, Rosemarie Chenard-Soucy^5^, Tommy Chevrette^5^, Emmanuelle Cloutier^5^, Richard Cloutier^5^, Hugues Cormier^5^, Gilles Cote^5^, Joanne Cyr^5^, Pierre David^5^, Luigi De Benedictis^5^, Marie-Claude Delisle^5^, Patricia Deschenes^5^, Cindy D. Desjardins^5^, Gilbert Desmarais^5^, Jean-Luc Dubreucq^5^, Mimi Dumont^5^, Alexandre Dumais^5^, Guylaine Ethier^5^, Carole Feltrin^5^, Amelie Felx^5^, Helen Findlay^5^, Linda Fortier^5^, Denise Fortin^5^, Leo Fortin^5^, Nathe Francois^5^, Valerie Gagne^5^, Marie-Pierre Gagnon^5^, Marie-Claude Gignac-Hens^5^, Charles-Edouard Giguere^5^, Roger Godbout^5^, Christine Grou^5^, Stephane Guay^5^, Francois Guillem^5^, Najia Hachimi-Idrissi^5^, Christophe Herry^5^, Sheilah Hodgins^5^, Saffron Homayoun^5^, Boutheina Jemel^5^, Christian Joyal^5^, Edouard Kouassi^5^, Real Labelle^5^, Denis Lafortune^5^, Michel Lahaie^5^, Souad Lahlafi^5^, Pierre Lalonde^5^, Pierre Landry^5^, Veronique Lapaige^5^, Guylaine Larocque^5^, Caroline Larue^5^, Marc Lavoie^5^, Jean-Jacques Leclerc^5^, Tania Lecomte^5^, Cecile Lecours^5^, Louise Leduc^5^, Marie-France Lelan^5^, Andre Lemieux^5^, Alain Lesage^5^, Andree Letarte^5^, Jean Lepage^5^, Alain Levesque^5^, Olivier Lipp^5^, David Luck^5^, Sonia Lupien^5^, Felix-Antoine Lusignan^5^, Richard Lusignan^5^, Andre J. Luyet^5^, Alykhanhthi Lynhiavu^5^, Jean-Pierre Melun^5^, Celine Morin^5^, Luc Nicole^5^, Francois Noel^5^, Louise Normandeau^5^, Kieron O’Connor^5^, Christine Ouellette^5^, Veronique Parent^5^, Marie-Helene Parizeau^5^, Jean-Francois Pelletier^5^, Julie Pelletier^5^, Marc Pelletier^5^, Pierrich Plusquellec^5^, Diane Poirier^5^, Stephane Potvin^5^, Guylaine Prevost^5^, Marie-Josee Prevost^5^, Pierre Racicot^5^, Marie-France Racine-Gagne^5^, Patrice Renaud^5^, Nicole Ricard^5^, Sylvie Rivet^5^, Michel Rolland^5^, Marc Sasseville^5^, Gabriel Safadi^5^, Sandra Smith^5^, Nicole Smolla^5^, Emmanuel Stip^5^, Jakob Teitelbaum^5^, Alfred Thibault^5^, Lucie Thibault^5^, Stephanye Thibault^5^, Frederic Thomas^5^, Christo Todorov^5^, Valerie Tourjman^5^, Constantin Tranulis^5^, Sonia Trudeau^5^, Gilles Trudel^5^, Nathalie Vacri^5^, Luc Valiquette^5^, Claude Vanier^5^, Kathe Villeneuve^5^, Marie Villeneuve^5^, Philippe Vincent^5^, Marcel Wolfe^5^, Lan Xiong^5^, Angela Zizzi^5^

---

**Supplementary Figure 1.**
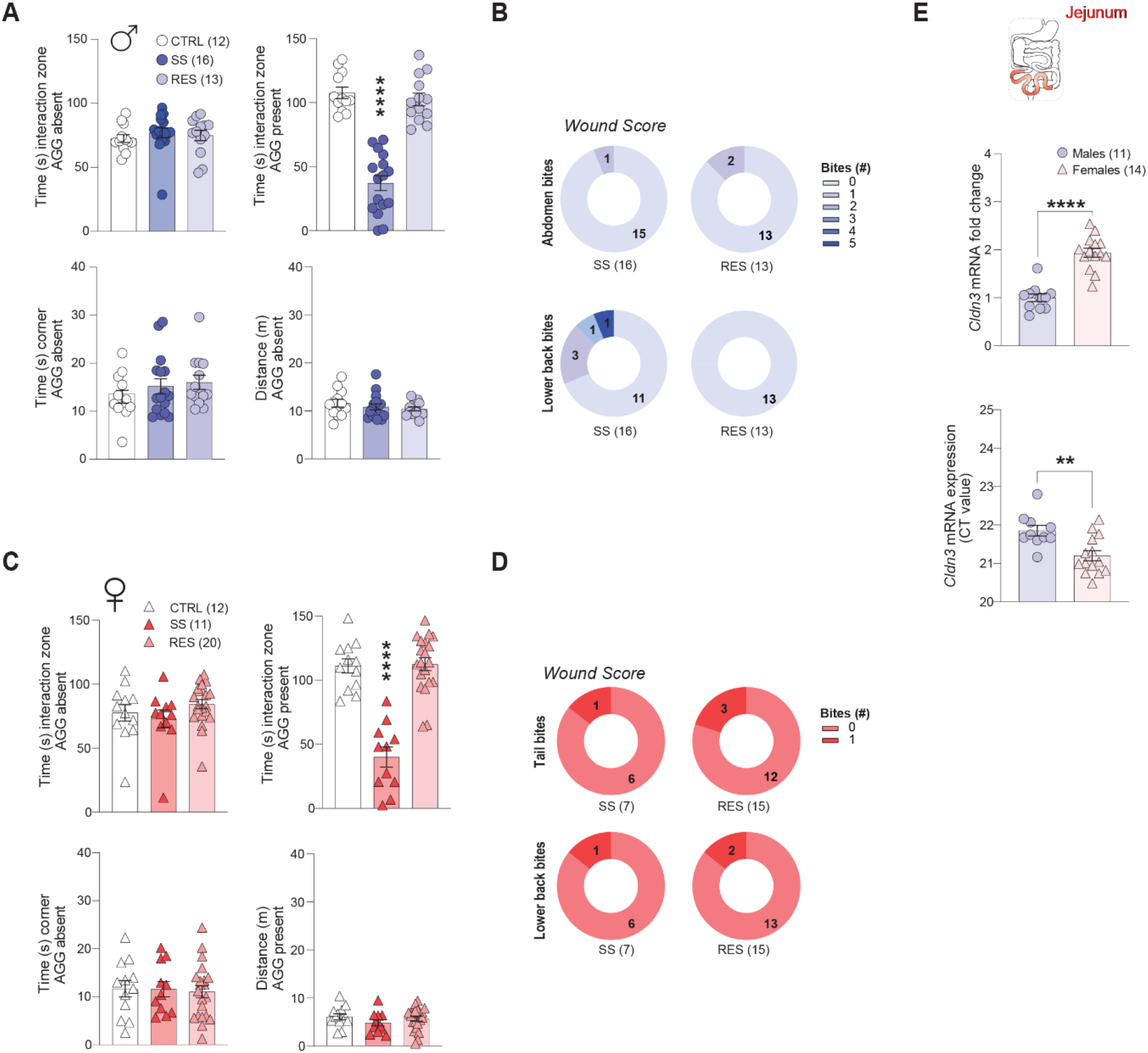
Behavioral phenotyping of quantitative PCR experiments and jejunum *Cldn3* expression is higher in female at baseline. **A)** Stress-susceptible (SS) male mice spent less time in the interaction zone when the social target (aggressor, AGG) was present when compared to unstressed controls (CTRL) and resilient (RES) animals. No significant difference was measured for time spent in the interaction zone or in the corners when the social target is absent. No difference was observed for locomotion. **B)** Wounding was comparable between stressed males. **C)** SS female mice spent less time in the interaction zone when the AGG was present when compared to CTRL and RES animals. No significant difference was measured for locomotion, time spent in the interaction zone or in the corners when the social target is absent. **D)** Wounding was comparable between stressed females. **E)** *Cldn3* mRNA level in the jejunum is higher in control females if compared to males from the same group. Data are assessed by two-tailed T-tests and one-way ANOVA followed by Bonferroni’s multiple comparison test for changes between groups; \*\**p*<0.01, \*\*\*\**p*<0.0001.

**Supplementary Figure 2.**
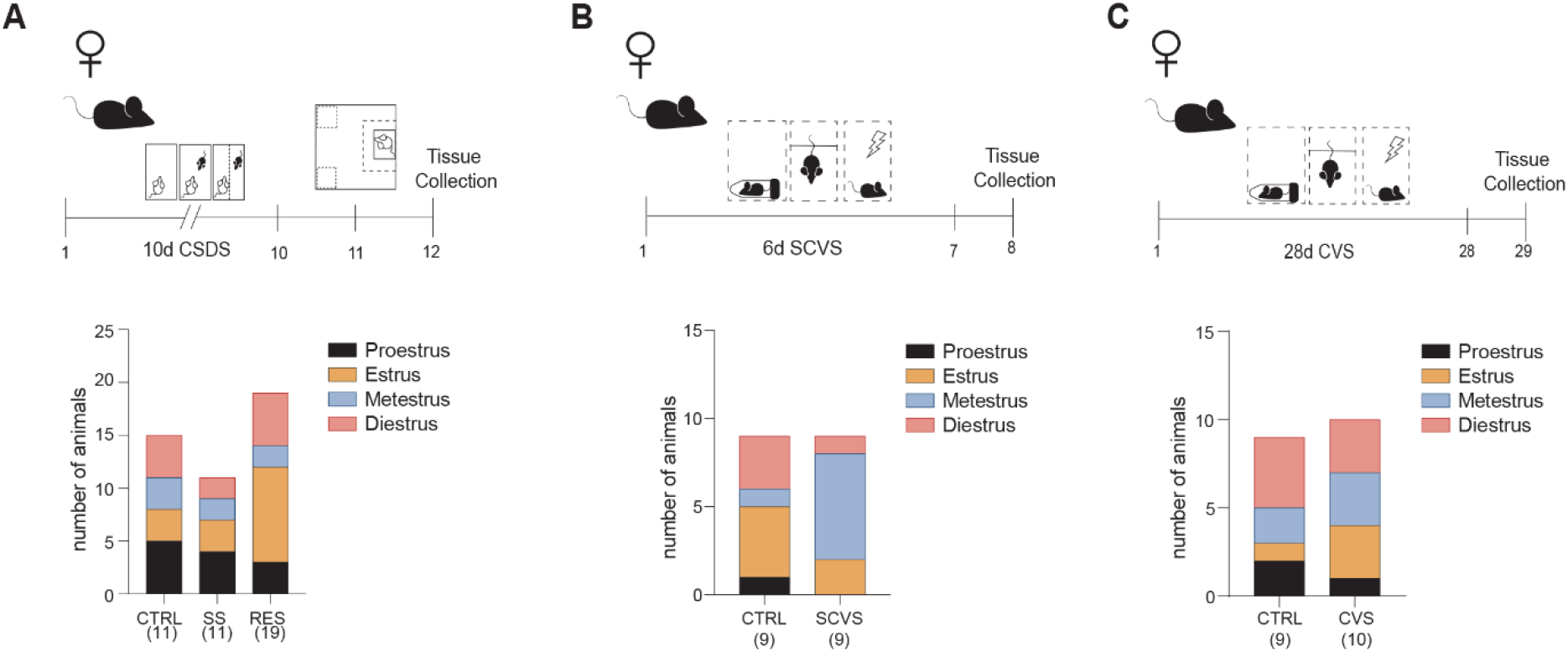
Phenotype vs phase of the estrus cycle of female mice. No significant difference was observed for the estrus cycle of female mice exposed to 10-d chronic social defeat stress (**A**, CSDS), 6-d subchronic variable stress (**B**, SCVS) or 28-d chronic variable stress (**C**, CVS).

**Supplementary Figure 3.**
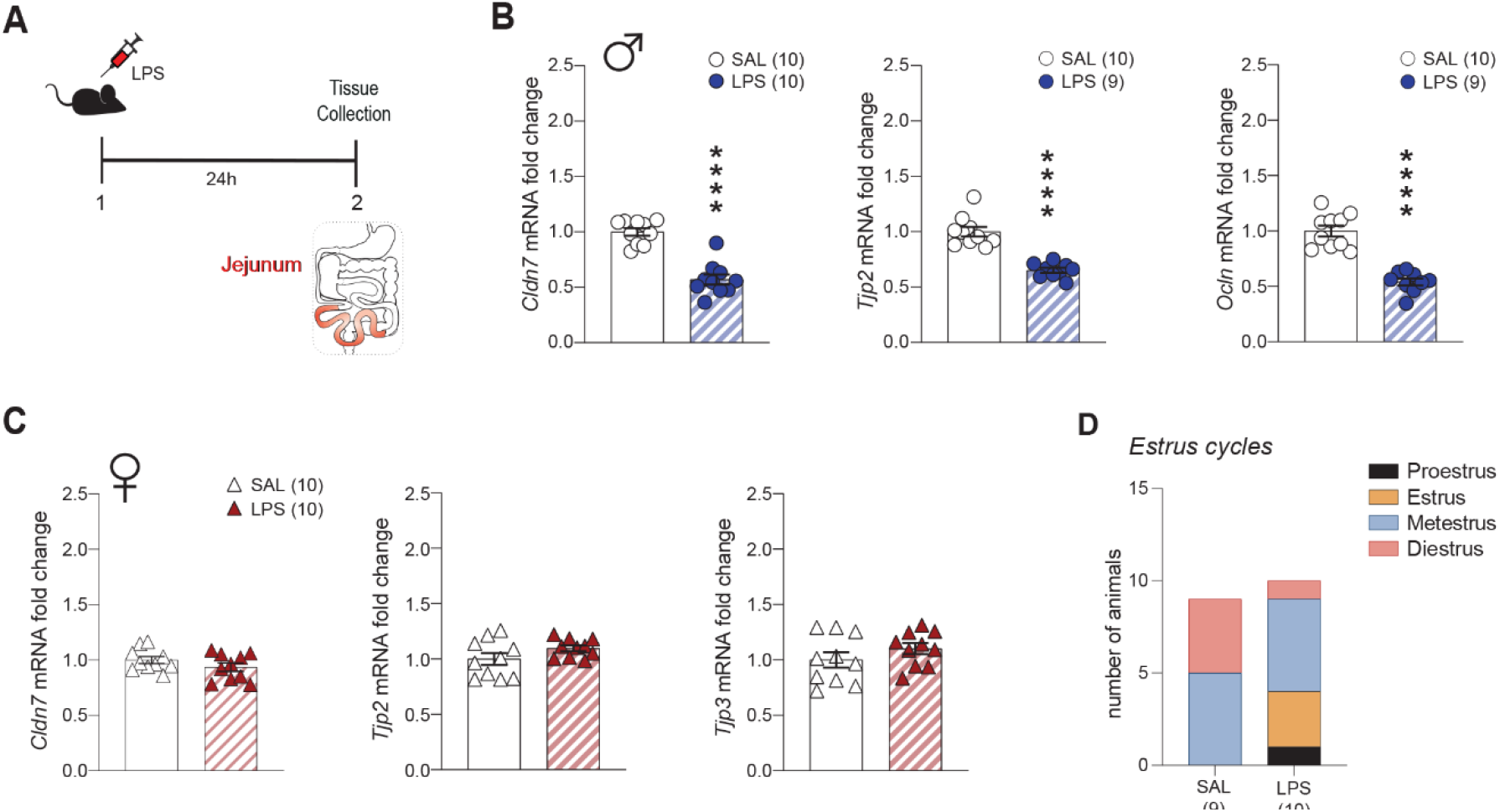
Additional jejunum tight junction gene expression data following LPS injection in male and female mice. **A)** Experimental timeline for pharmacological treatment with LPS and tissue collection. **B)** Gut tight junction-related Claudin-7 (*Cldn7*), Tight junction protein 2 (*Tjp2*) and Occludin (*Ocln*) are decreased in the jejunum (JEJ) of male mice following LPS injection. **C)** Conversely, no change was observed for JEJ *Cldn7, Tjp2* or *Tjp3* expression for females despite exposure to the same treatment. **D)** Estrus cycle phase was similar between groups for female injected or not with LPS. Two-tailed t-test was performed to evaluate changes between groups; \*\*\*\**p*<0.0001.

**Supplementary Figure 4.**
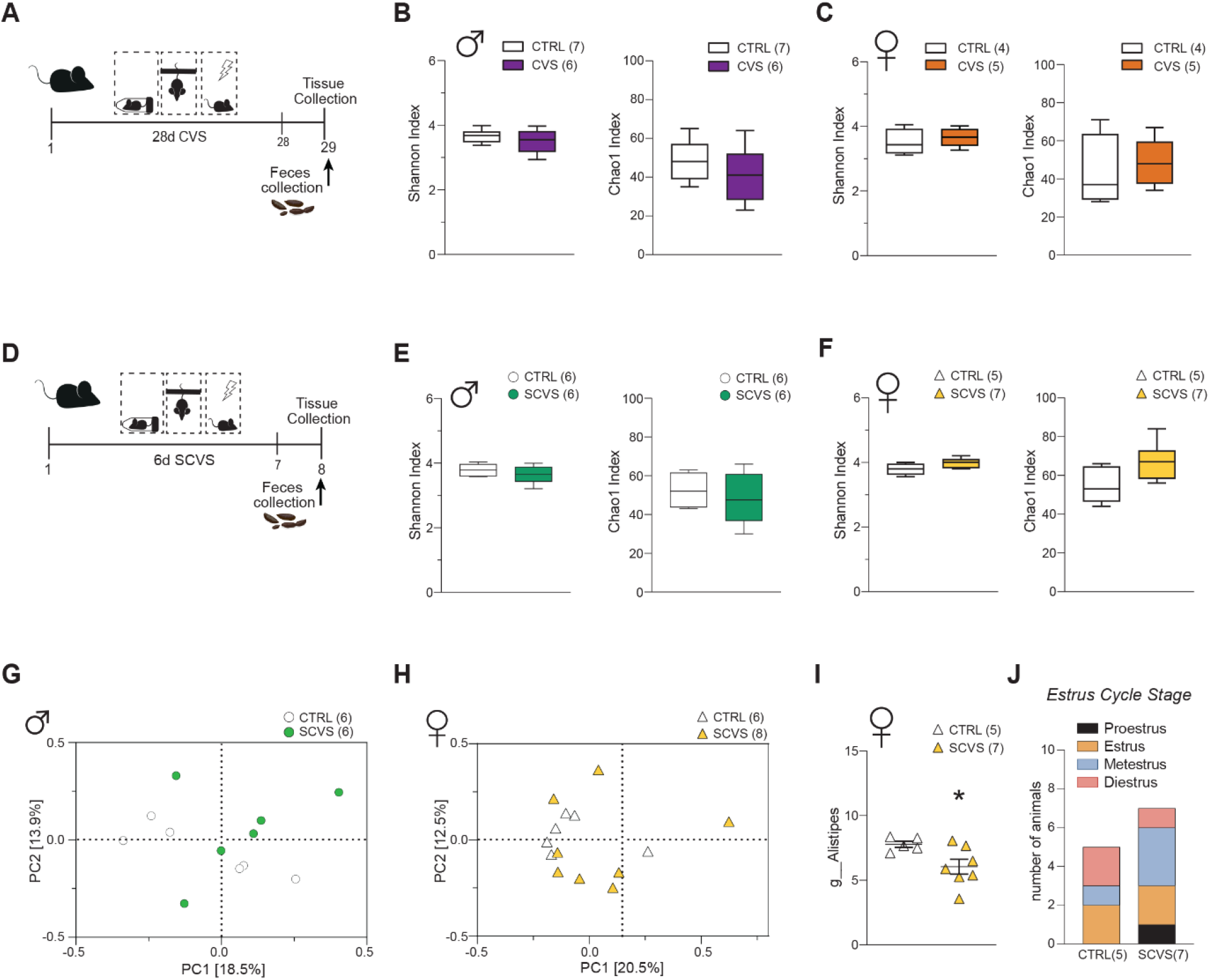
Additional microbiota-related data following chronic and subchronic variable stress paradigms in male and female mice. **A)** Experimental timeline of feces collection after 28-d chronic variable stress (CVS). No significant difference was noted for the Shannon or Chao1 alpha-diversity indices for males (**B**) or females (**C**) after CVS exposure. **D)** Experimental timeline of feces collection after 6-d subchronic variable stress (SCVS). **E)** Again, no significant effect was observed for males (**E, G**) or female (**F, H**) Shannon or Chao1 alpha-diversity indices or beta-diversity. **I)** As for females, only *Alistipes* was reduced (*p=0.0370*) despite exposure to the same stressor. **J)** Estrus cycle did not have an impact. Unpaired t-tests were performed for two-group comparisons; \**p*<0.05, \*\**p*<0.01.

**Supplementary Figure 5.**
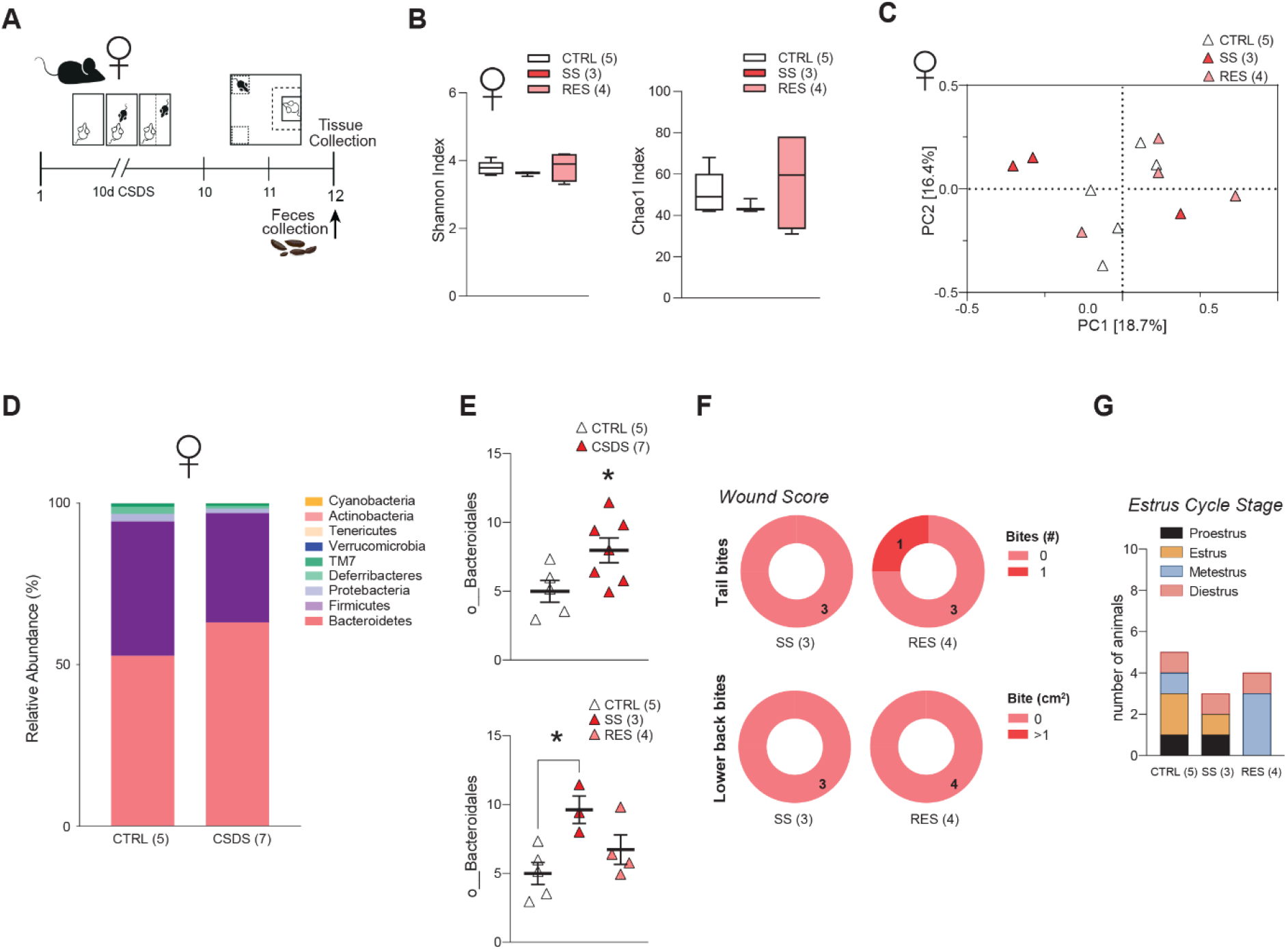
Additional microbiota-related data and phenotyping for female chronic social defeat stress. **A)** Experimental timeline of feces collection after 10-d chronic social defeat stress (CSDS) exposure. **B)** No difference was noted between unstressed controls (CTRL), stress-susceptible (SS) or resilient (RES) female mice for the alpha-diversity Shannon or Chao1 indices or beta-diversity (**C**). **D)** Microbiota phyla relative abundance did not reveal significant differences between unstressed controls or mice subjected to CSDS. **E)** However, an increase in unassigned members of *Bacteroidales* was apparent in the stressed vs CTRL group (*p=0.0411*) and this effect was driven by the SS females (*p=0.0270*). **F)** Wound scores were comparable between stressed female mice as assessed by the number of tail bites and lower back bites. **G)** Estrus cycle phase was also similar. One-way ANOVA or unpaired t-test was performed for three- or two-group comparisons, respectively; \**p*<0.05.

**Supplementary Figure 6.**
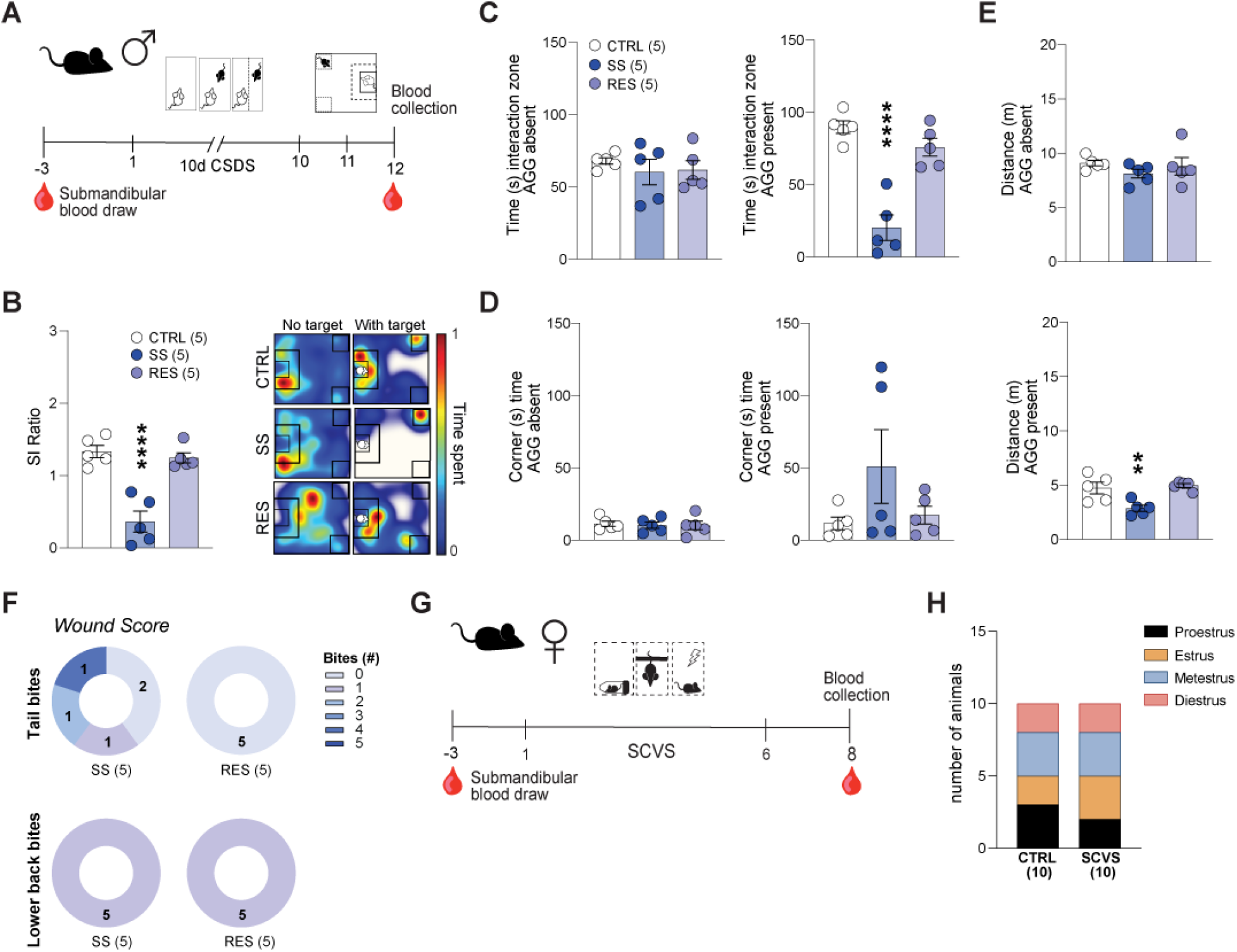
Behavioral phenotyping for blood-based analysis of gut-related biomarkers. **A)** Experimental timeline of 10-d chronic social defeat stress (CSDS) and blood collection prior and after stress exposure. Stress-susceptible (SS) mice had lower social interaction (SI) ratio (**B**) since they spent less time in the interaction zone when the social target (aggressor, AGG) was present (**C**). No difference was observed for time spent in the corners (**D**) or for locomotion when the AGG is absent (**E**). **F)** Wound scores were comparable between stressed male mice as assessed by the number of tail bites and lower back bites. **G)** Experimental timeline of 6-d subchronic variable stress (SCVS) and blood collection prior and after stress exposure. **H)** No significant difference was noted for estrus cycle phase of female mice subjected or not to 6-d SCVS. One-way ANOVA followed by Bonferroni’s multiple comparison test for changes between groups; \*\**p*<0.01, \*\*\*\**p*<0.0001.

**Supplementary Table 1.** Quantitative PCR primers.

| Gene | Assay ID |
| --- | --- |
| **Cldn3** | Mm.PT.58.43310459.g |
| **Cldn7** | Mm.PT.58.16298059 |
| **Cldn12** | Mm.PT.58.41535303 |
| **Tjp1** | Mm.PT.58.29459730 |
| **Tjp2** | Mm.PT.58.16834535 |
| **Tjp3** | Mm.PT.58.43961106 |
| **Ocln** | Mm.PT.58.42749240 |
| **Marveld2** | Mm.PT.58.7719303 |
| **Muc2** | Mm.PT.58.29496069.g |
| **Ido1** | Mm.PT.58.29540170 |
| **Ahr** | Mm.PT.58.11116644 |
| **Actb** | Mm.PT.39a.22214843.g |

**Supplementary Table 2.**
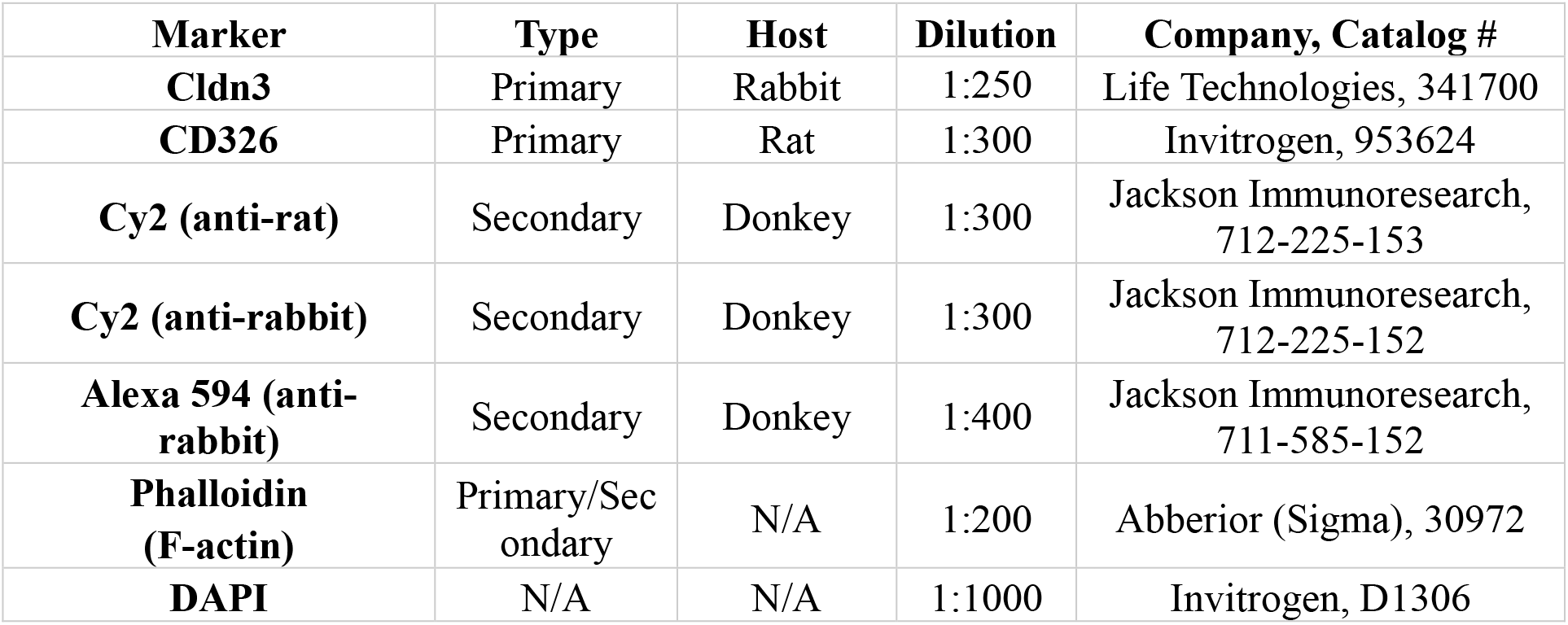
Primary and secondary antibodies.

**Supplementary Table 3.**
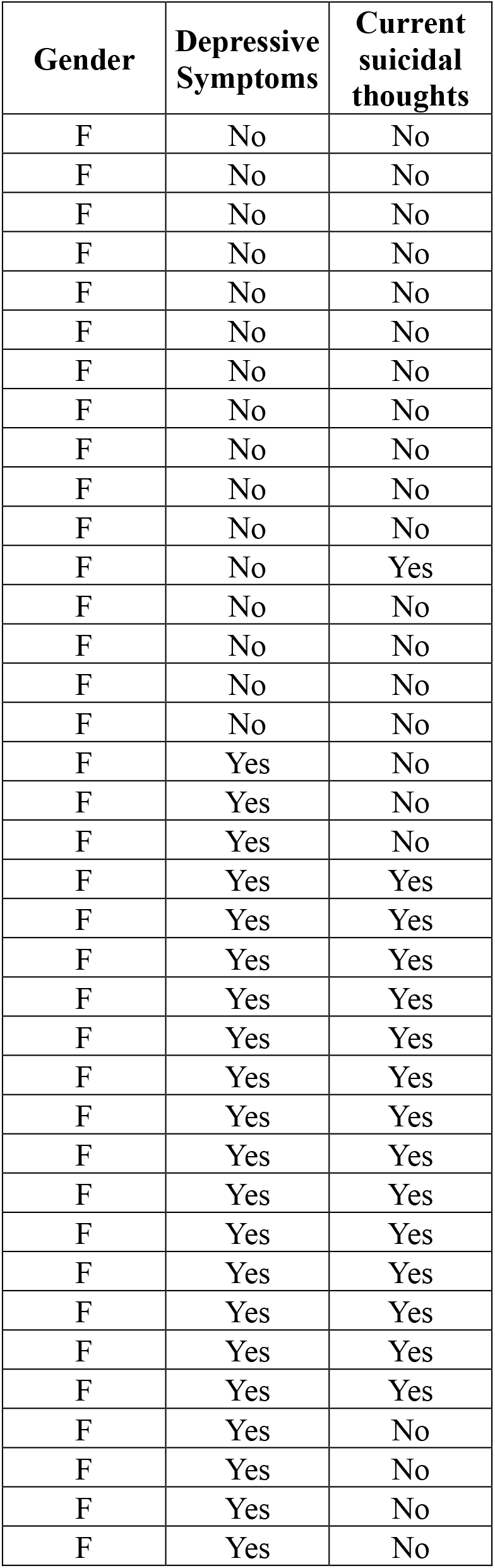
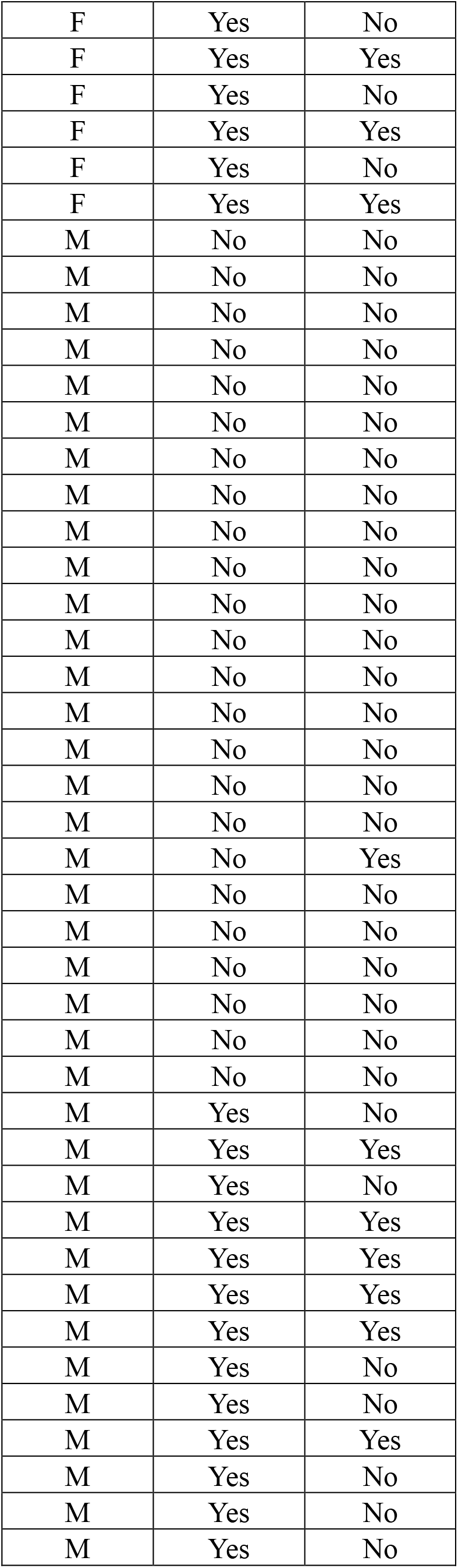
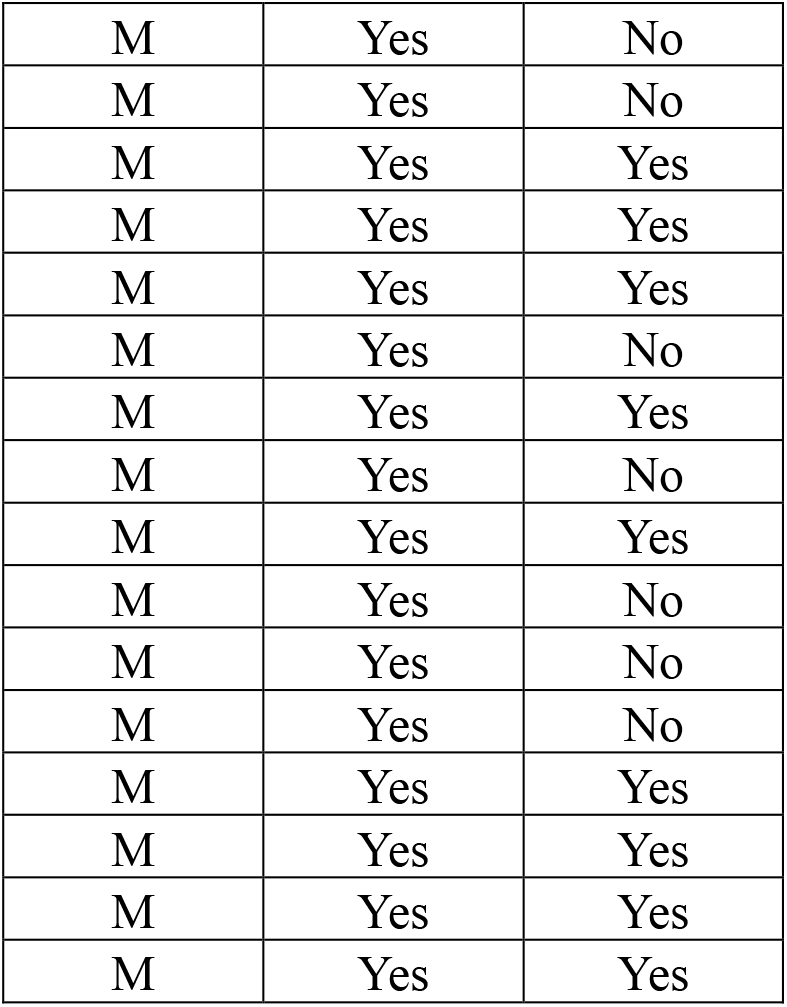
Complete demographic data for human serum cohort. Age of the participants (years): 18-30

## Notes

### Competing Interest Statement

The authors have declared no competing interest.

